# Flap endonuclease 1 repairs DNA-protein crosslinks via ADP-ribosylation

**DOI:** 10.1101/2023.10.19.563118

**Authors:** Yilun Sun, Lisa M. Jenkins, Lara H. El Touny, Ukhyun Jo, Xi Yang, Tapan K. Maity, Liton K. Saha, Isabel Uribe, Sourav Saha, Shunichi Takeda, Anthony K.L. Leung, Ken Cheng, Yves Pommier

## Abstract

DNA-protein crosslinks (DPCs) are among the most ubiquitous and detrimental DNA lesions which arise from exposure to metabolic stresses, drugs, or crosslinking agents such as formaldehyde (FA). FA is a cellular by-product of methanol metabolism, histone demethylation, lipid peroxidation as well as environmental pollutants. Failure to repair FA-induced DPCs blocks nearly all chromatin-based processes including replication and transcription, leading to immunodeficiencies, neurodegeneration, and cancer. Yet, it remains largely unknown how the cell repairs DPCs. The study of DPC repair is impeded by our incomprehension of the types of proteins crosslinked by FA due to the lack of techniques to identify the DPCs. Here, we designed a novel bioassay to profile FA-induced DPCs by coupling cesium chloride differential ultracentrifugation with HPLC-mass spectrometry (MS). Using the method, we revealed the proteome of FA-induced DPCs in human cells and found that the most abundant proteins that form DPCs are PARP1, topoisomerases I and II, methyltransferases, DNA and RNA polymerases, histones, as well as ribosomal proteins. To identify enzymes that repair DPCs, we carried out RNA interference screening and found that downregulation of flap endonuclease 1 (FEN1) rendered cells hypersensitive to FA. Since FEN1 possesses 5’-flap endonuclease activity, we hypothesized that FA induces DPC-conjugated 5’-flap DNA fragments that can be processed by FEN1. Indeed, we demonstrate that FA damages DNA bases that are converted into 5’-flap via the base excision pathway (BER). We also observed that the damaged DNA bases were colocalized with DPCs and FEN1. Mechanistically, we showed that FEN1 repairs FA-induced DPCs *in vivo* and cleaves 5’-flap DNA substrate harboring DPC mimetic *in vitro*. We also found that FEN1 repairs enzymatic topoisomerase II (TOP2)-DPCs induced by their inhibitors etoposide and doxorubicin independently of the BER pathway, and that FEN1 and the DPC-targeting protease SPRTN act as parallel pathways for the repair of both FA-induced non-enzymatic DPCs and etoposide-induced enzymatic TOP2-DPCs. Notably, we found that FA-induced non-enzymatic DPCs and enzymatic TOP2-DPCs are promptly modified by poly-ADP-ribosylation (PARylation), a post-translational modification catalyzed by PARP1, a key DNA damage response effector that acts by PARylating both DNA damage sites and DNA repair proteins. We performed immunoprecipitation (IP) assays with anti-PAR antibody for HPLC-MS and identified FEN1 as a PARylation substrate. Next, we showed that PARylation of DPC substrates signaled FEN1 whereas PARylation of FEN1 drove FEN1 to DPC sites. Finally, using the enzymatic labeling of the terminal ADP-ribose-MS method, we identified the E285 residue of FEN1 as a dominant PARylation site, which appeared to be required for FEN1 relocation to DPCs. Taken together, our work not only unveiled the identities of FA-induced DPCs but also discovered an unprecedented PARP1-FEN1 nuclease pathway as a universal and imperative mechanism to repair the miscellaneous DPCs and prevent DPC-induced genomic instability.

## Introduction

DNA-protein interactions are at the heart of normal cell function ^1^. In eukaryotic cells, genomic DNA is wrapped around histone octamers to allow for chromosomal packaging. Binding of specific proteins to DNA directs replication and chromosome remodeling, modulates transcription, and governs DNA damage responses. Because of their fundamental significance in all cellular processes involving DNA, dynamic DNA-protein interactions are required for cell survival, and their disruption has serious biological consequences^2^.

Chromatin-interacting proteins are susceptible to covalent trapping on DNA strands upon exposure to endogenous lesions or exogenous (environmental and chemotherapeutic) agents ^3–6^, leading to the formation of DNA-protein crosslinks (DPCs). According to their chemical nature, DPCs can be divided into two classes. First, intermediate DNA-protein covalent conjugates that arise as part of normal enzymatic reaction (e.g. topoisomerase, DNA polymerase, and DNA methyltransferase intermediates) can be trapped under ubiquitous circumstances including enzyme malfunctioning, exposure to endogenous DNA damage, and treatments with their respective inhibitors ^7, 8^. Second, non-enzymatic DPCs are caused by non-specific crosslinking of proteins in the vicinity of DNA, which can be caused by endogenous reactive metabolites as well as exogenous agents. As a result of endogenous exposure to reactive oxygen species, lipid peroxidation products, as well as normal cellular metabolism, non-enzymatic DPCs progressively accumulate in the brain and many other tissues ^9^. Because of their considerable size and their helix-distorting nature, DPCs interfere with the progression of replication and transcription machineries and hence hamper the faithful expression of genetic information. DPC-generated DNA damage induces cancer, neurodegeneration, immunodeficiency, hepatoxicity, and premature aging ^5, 10–12^.

An important endogenous source of non-enzymatic DPCs is aldehydes. Reactive formaldehyde (FA) crosslinks proteins to DNA via a methylene group that bridges the exocyclic amine of DNA bases and protein amino acids (most commonly a lysine L-amino group) to form a secondary amine (-NH-CH-NH-). FA is generated as a byproduct of different metabolic processes such as methanol metabolism, histone demethylation, or lipid peroxidation; it is also an important environmental pollutant ^5^. FA is found at high concentrations in human plasma and certain tissues, suggesting that DPCs can be continuously induced by FA and cells must constantly overcome DPC-induced damage.

The repair of DPCs has been extensively studied over the past two decades, especially for the enzymatic DPCs generated by DNA topoisomerases. It involves complex mechanisms encompassing multiple repair proteins ^4, 6, 9^. While the overall pathway(s) of DPC repair has yet to be fully unraveled, a large body of recent evidence has demonstrated that proteolysis of the bulky protein component plays a pivotal role in the repair of DPCs including non-enzymatic DPCs. Proteolysis can be achieved by different proteases under different cellular contexts. The DNA-dependent metalloprotease SPRTN (Wss1 in yeast) acts on enzymatic and non-enzymatic DPCs during replication as a component of the replisome^13–15^ whereas the 26S proteasome can digest DPCs both in and outside of S-phase^16–20^. In addition to SPRTN, several proteases, such as ACRC^21^, also known as germ cell nuclear antigen (GCNA)^22^, FAM111A and FAM111B^23, 24^, and DDI1 and DDI2^25, 26^, have recently been discovered and linked to DPC proteolysis.

Yet, the methylene groups formed by FA link an amino group of a DNA base and a lysine L-amino group of a protein (secondary amine) and are not substrates for peptidases. Therefore, none of the known proteases can fully excise the FA-induced DPCs, and it can be speculated that cells employ a nucleolytic process to remove the DPCs. To discover such repair nucleases, we first developed a mass spectrum-based method to identify the proteins that form DPCs upon exposure to FA. This approach led to the identification of proteins that form enzymatic DPCs and DPC-like trapping (TOP1, TOP2α, TOP2β, DNMT1, PARP1) as some of the most abundant DPCs induced by FA. Next, we performed an unbiased RNA interference (RNAi) screen with the Horizon human DNA damage response library and found several DNA repair nucleases mitigating FA-induced cytotoxicity. Using the modified RADAR (rapid approach to DNA adduct recovery) assay ^17^ to measure FA-induced DPC *in vivo*, we identified flap endonuclease 1 (FEN1) as a critical nuclease for non-enzymatic DPC resolution in parallel to the SPRTN protease pathway. Flaps are single-stranded DNA structures that can arise when the replication machinery encounters damaged DNA. FEN1 cleaves 5’-flap structures to generate a nick in the DNA strand as an endonuclease, allowing the replication or repair machinery to continue its work ^27^. As FEN1 is a replication-associated nuclease involved in the prominent base excision repair (BER) and Okazaki fragment maturation pathways ^28, 29^, we hypothesized that FEN1 excises DPCs formed within a DNA flap generated by the BER pathway and formed within the RNA-DNA hybrid flaps of Okazaki fragments. We show that FA-induced oxidative lesions co-localized with DPCs and FEN1, in line with an earlier study reporting that FEN1 prevents the formation of 2-deoxyribonolactone (an oxidized abasic site)-induced DPC formation^30^. We also validate the activity of FEN1 against DNA flap harboring amino acid mimetic *in vitro*.

Notably, we show that FEN1 also repairs the enzymatic topoisomerase II (TOP2)-DPC. The TOP2 homodimer breaks both DNA strands and forms a covalent enzyme-DNA intermediate termed TOP2 cleavage complex or TOP2cc, which consists of a 5′ phosphotyrosyl linkage between the enzyme and DNA^31^. The transient TOP2cc is typically reversed once the enzyme reseals the breakage following strand passage but can be converted into a long-lived TOP2-DPCs under certain conditions including exposure to its inhibitors. While the TOP2-DPCs can be repaired by the proteases including SPRTN as well as by TDP2, a specialized nuclease that hydrolyzes the phosphotyrosyl linkage^19, 20^, we found that FEN1 provides a salvage route to repair the 5’ enzymatic TOP2α- and β (two isozymes in mammals)-DPCs with its endonuclease activity independently of the SPRTN and TDP2 pathways. Unlike the repair of FA-induced nonenzymatic DPCs, FEN1 appears to resolve the enzymatic TOP2-DPCs without the help of the BER pathway.

Considering that PARylation plays a central role in sensing DNA damage and that we found that PARP1-DPCs were among the most abundant DPCs induced by FA, we conjectured that PARylation participates in DPC repair by regulating FEN1. Accordingly, we observed PARylation of FA-induced non-enzymatic DPCs and etoposide-induced enzymatic TOP2-DPCs by blocking PARG (PAR glycohydrolase) ^32^. We also found that FEN1 is a substrate of PARylation, which is stimulated upon FA exposure. We profiled and identified a specific FEN1 PARylation site E285 amino acid residue crucial for the recruitment of FEN1 to the DPC site. In summary, our study not only identifies the proteins involved in the DPCs generated by FA but also an unprecedented FEN1 pathway that excises both non-enzymatic and enzymatic DPCs via PARP1-mediated ADP-ribosylation, shedding new light on the repair of DPCs and the etiology of DPC-associated diseases.

## Result

### 1. Identification of formaldehyde (FA)-induced DNA-protein crosslink (DPC) proteome by coupling in vivo complex of enzyme bioassay and quantitative mass spectrometry

Pathway choice for DNA damage repair is largely contingent on cellular contexts (replication, transcription, chromosome remodeling, and segregation, etc.) ^33^. Given the heterogeneity of the non-enzymatic DPCs induced by FA, the repair of those DPCs remains only partially known. Understanding the protein identity of as well as chromosome regions, DNA transactions and DNA structures involved in the DPCs is therefore important to study ^34, 35^ non-enzymatic DPC repair. A major challenge to profiling non-enzymatic DPCs is their purification.

To solve this, we isolated FA-induced DPC-containing genomic DNA using cesium chloride-based ultracentrifugation, a method derived from the *in vivo* complex of enzyme (ICE) assay, followed by release of the crosslinked proteins using micrococcal nuclease ^17^ and profiling of the liberated proteins by mass spectrometry analysis (Fig. 1A). Based on this ICE-MS method, we unveiled the proteome of FA-induced DPCs in three human cell lines (U2OS, HEK293, and MCF7) and found that the most abundant proteins that form DPCs are PARP1, topoisomerases I and II (TOP1 and 2), methyltransferases including DNMT1, DNA and RNA polymerases, histones, as well as ribosomal proteins (Fig. 1B-D; Supplementary fig. 1), of which TOP1, TOP2, DNMT as well as PARP1 are also known to form enzymatic DPCs after exposure to endogenous DNA lesions and their inhibitors. We confirmed this finding by conducting a modified Rapid Approach to DNA Adduct Recovery (RADAR) assay ^34–36^ to immunodetect these DPCs in cells and observed induction of TOP1-, TOP2α- and β-, DNMT1-, PARP1-, as well as RNA Pol II-DPCs by FA in a dose-dependent manner (Fig. 1E and F). Considering the involvement of TOP1, TOP2, PARP1, and DNMT1 in the DNA replication ^8, 37, 38^, we hypothesized that the repair of these DPCs by a nucleolytic mechanism is likely coupled with active replication.

**Figure 1.**
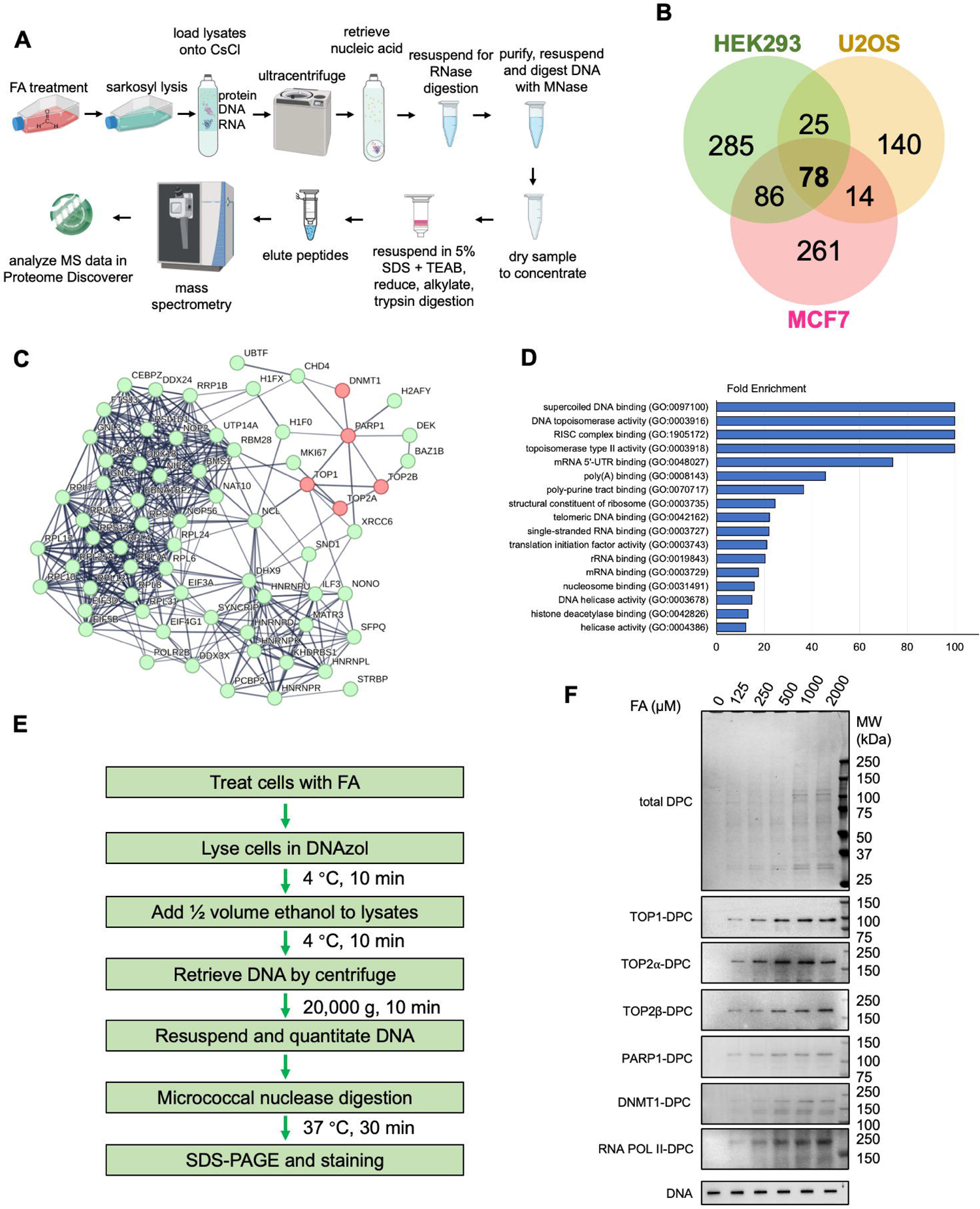
Identification of the FA-induced DPC proteome by ICE-MS. **A.** Experimental scheme of ICE-MS profiling of DPCs induced by FA. **B.** Venn diagrams depicting the number of DNA-crosslinked proteins significantly upregulated by FA (10-fold increase over control) across the indicated three cell lines. **C.** Common DNA-crosslinked proteins upregulated by FA across the three cell lines were subjected to network analysis using the STRING database, at the interaction confidence setting of 0.7. Proteins not connected to the network were omitted. Proteins that form enzymatic DPCs are highlighted in light red. **D.** The common FA-induced protein adducts across the three cell lines were mapped to the human proteome, which was annotated with Gene Ontology (GO) molecular function. Enrichment analysis was performed by fold change measurement. **E.** Experimental scheme of the modified RADAR assay for DPC detection. **F.** The modified RADAR assay was performed in HEK293 cells treated with FA at the indicated concentrations for 2 hours. Ten µg samples were digested with 100 units micrococcal nuclease and subjected to SDS-PAGE electrophoresis. Total DPCs were detected with Coomassie stain and individual DPCs were probed with antibodies targeting the corresponding proteins. 2 µg samples without micrococcal nuclease digestion were subjected to slot-blot and probed with anti-DNA antibody as loading control.

### 2. Flap endonuclease 1 (FEN1) is a major nucleolytic mechanism for non-enzymatic DPC repair

To identify the nuclease(s) required for non-enzymatic DPC repair, we conducted an RNA interference (RNAi) high throughput screen using a human ON-TARGETplus short-interfering RNA (siRNA) library targeting 240 genes involved in the DNA damage and repair pathways including nearly all the DNA repair nucleases. Following transfection of a set of 4 individual siRNAs for each gene in MCF7 cells in a 384-well plate format, we exposed the cells to 80 µM FA for 72 hours and subsequently measured their survival using the CellTiter-Glo luminescent assay (Fig. 2A). FA responses were converted to z-scores from the 4-individual siRNA-transfected wells. The normalized z-score data were plotted for each gene, and negative median z-scores were considered synthetic-lethal interactions with FA treatment. ERCC3, ERCC6, and DNMT1 were identified as the most potent hits. MRE11, ERCC4, MUS81, FEN1, ERCC5, EXO1, APEX1, SLX1A, and DNA2 nucleases were also identified as synthetic lethal with FA (Fig. 2B).

**Figure 2.**
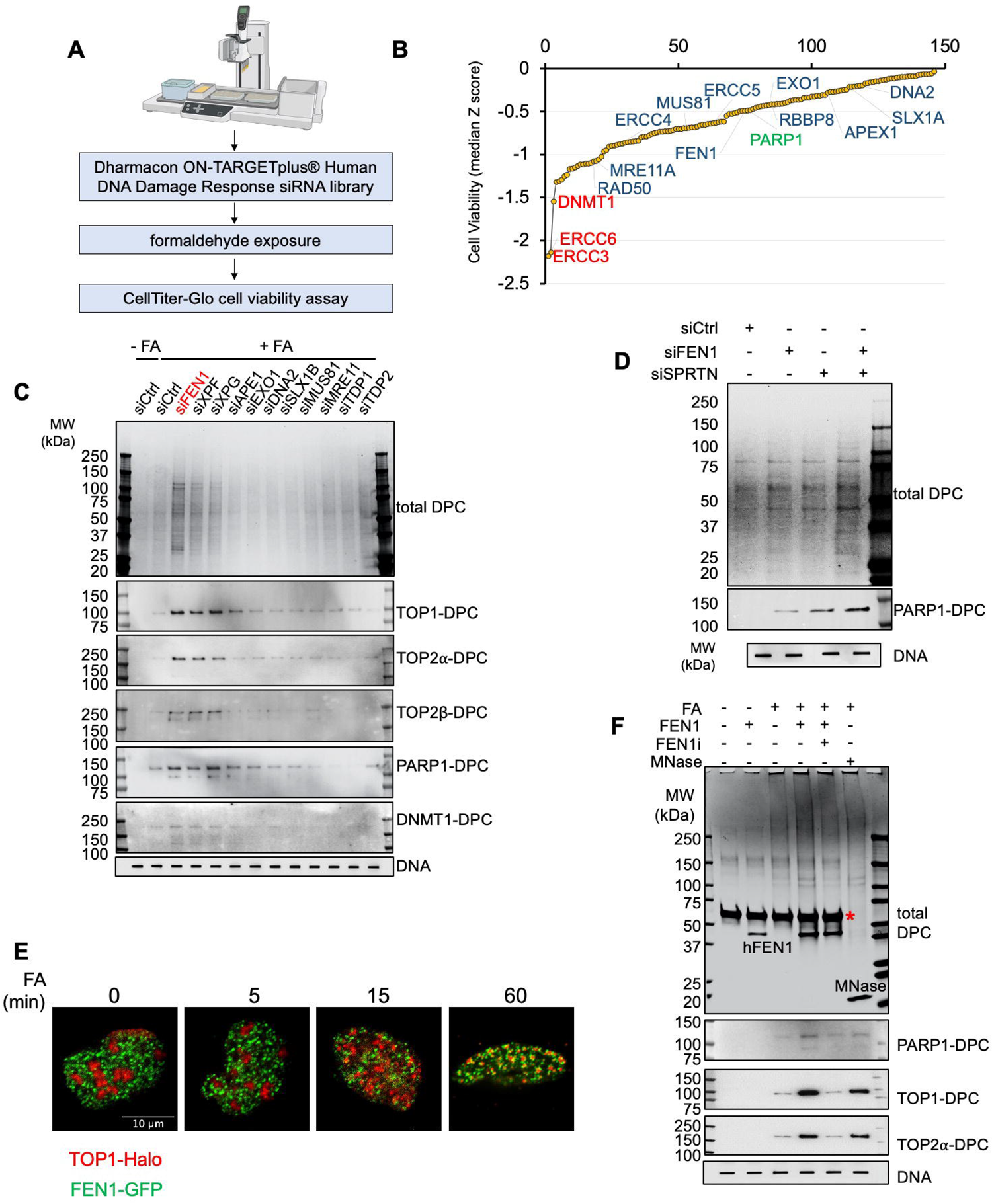
FEN1 is a major nucleolytic mechanism for non-enzymatic DPC repair. **A.** Schematic overview of the RNAi screen workflow in MCF7 cells. **B.** Ranked distribution plots of median Z-scores obtained from RNAi screen in FA-treated/untreated MCF7 cells. Hit genes encoding DNA repair nucleases selected for further validation are highlighted in blue. The top 3 sensitizing genes are highlighted in red. **C.** The modified RADAR assay was performed in HEK293 cells transfected with indicated siRNAs and subsequently treated with 400 µM FA for 4 hours. Total DPCs were detected with Coomassie stain and PARP1-DPCs were probed withanti-PARP1 antibody. **D.** The modified RADAR assay was performed in HEK293 cells transfected with indicated siRNAs and subsequently treated with 400 µM FA for 2 hours. Total DPCs were detected with Coomassie stain and individual DPCs were probed with antibodies targeting the indicated proteins. **E.** Instant structure illumination microscopy analysis in live U2OS cells transfected with TOP1-HaloTag and FEN1-GFP constructs and subsequently exposed to 400 µM FA for indicated periods of time. **F.** The modified RADAR assay was performed in HEK293 cells treated with or without FA (400 µM, 2 h). Samples were incubated with 500 nM recombinant human FEN1 in FEN1 cleavage buffer or with 100 units micrococcal nuclease in micrococcal nuclease cleavage buffer at 37°C for 30 min prior to SDS-PAGE electrophoresis. Recombinant human FEN1 was pre-incubated with or without 10 µM FEN1 inhibitor. *, Bovine serum albumin (BSA), stabilizer in FEN1 cleavage buffer.

To determine which nuclease helps cells to survive FA by facilitating the removal of FA-generated DPCs, we employed the modified RADAR assay to measure FA-induced DPC levels in cells transfected with siRNAs targeting the nucleases identified from the RNAi screen. Notably, we found that the downregulation of FEN1 markedly increased the levels of DPCs as detected by Coomassie blue and antibodies against individual DPCs (Fig. 2C; Supplementary fig. 2A). To confirm this result, we generated stable FEN1 knocked-down MCF7 cells with small hairpin RNA (shRNA) and carried out cell viability assays and the modified RADAR assays in the FEN1 KD cells treated with FA (Supplementary fig. 2B-D). Consistent with our hypothesis, we observed hypersensitivity to FA as well as accumulation of FA-induced DPCs in the FEN1-KD cells. We next carried out epistasis analysis for FEN1 and SPRTN, a well-established DPC-targeted protease. Double-knocking down (DKD) of FEN1 and SPRTN with their siRNAs exhibited an additional increase in the levels of FA-induced DPCs in comparison with their respective single knocking-down (Fig. 2D; Supplementary fig. 2E), suggesting FEN1 and SPRTN as redundant repair mechanisms for non-enzymatic DPC.

We next examined whether FEN1 and DPCs co-localize upon FA exposure. Based on reagent availability, we chose TOP1 as an example and monitored its subnuclear distribution in U2OS cells transfected with the TOP1-HaloTag expression construct. Prior to FA treatment, TOP1-HaloTag molecules appeared to be enriched in nucleoli. TOP1 proteins were rapidly delocalized from nucleoli to the nucleoplasm and co-localized with FEN1-GFP molecules following exposure to FA (Fig. 2E), which is consistent with prior findings that TOP1-DPCs induced by TOP1 inhibitors are expelled from nucleoli and localized to nucleoplasm ^39^.

Our finding that FEN1 and SPRTN serve as independent salvage routes for FA-induced DPC repair suggests that FEN1 resolves the DPCs without proteolysis of their protein components. To test this hypothesis, we incubated DNA samples isolated by the RADAR assay with recombinant human FEN1 instead of micrococcal nuclease and subjected the samples to SDS-PAGE. We found that the SDS gel separated recombinant FEN1-treated RADAR samples but not the samples without FEN1 treatment (Fig. 2F), as proteins crosslinked to megabase DNA molecules are trapped in the wells of the gel, suggesting that FEN1 acts on the full-length DPCs. Blocking FEN1 endonuclease activity with its small molecule inhibitor FEN1-IN-4 (FEN1i) ^40^ failed to release the RADAR samples for SDS-PAGE resolution (Fig. 2F), indicating that FEN1 processes DPCs using its 5’ overhanging flap endonuclease activity.

### 3. FEN1 cleaves DPC-harboring DNA flap substrates originated by oxidative lesions

As FEN1 only processes 5’-flap structures, and FA generates DPCs with no broken ends except DPCs formed close to the 5’-ends of Okazaki fragments in the lagging strand of newly synthesized DNA, we speculated that the no-break DPCs in the front of replication forks and on the leading strand behind the forks must be accompanied with 5’-DPC flaps in order for FEN1 to target and process these lesions (Fig. 3A). To test this hypothesis, we first examined whether FA induces base damage, as oxidative lesions can be converted to 5’-flaps by the BER pathway^41^. Accordingly, we observed that FA elevated cellular levels of 8-oxo-deoxyguanine (8-OXO-dG) (Fig. 3B), one of the major products of DNA oxidation, and cellular levels of AP sites (apurinic/apyrimidinic site) (Fig. 3C), which are formed by the processing of damaged bases by DNA glycosylase activity as the first step of the BER pathway. To further assess the role of the BER pathway in non-enzymatic DPC repair, we downregulated OGG1 (8-oxoguanine DNA glycosylase) and performed the modified RADAR assay. We found that OOG1 deficiency accumulated the DPCs but OGG1 and FEN1 DKD did not further increase the accumulation (Fig. 3D; Supplementary fig. 3A and B), suggesting that OGG1 likely acts upstream of FEN1 during non-enzymatic DPC repair. Next, we conducted the proximity ligation assay (PLA) and observed co-localization between TOP1 and 8-OXO-dG upon FA treatment (Fig. 3E; Supplementary fig. 3C). Because TOP1 acts in front of the replication fork to relax accumulated positive supercoils, this find suggests that FA generates adjoining damaged bases and DPCs ahead of the replication fork, which is likely transformed into flaps harboring the DPCs by the BER pathway, eventually leading to their resolution by FEN1.

**Figure 3.**
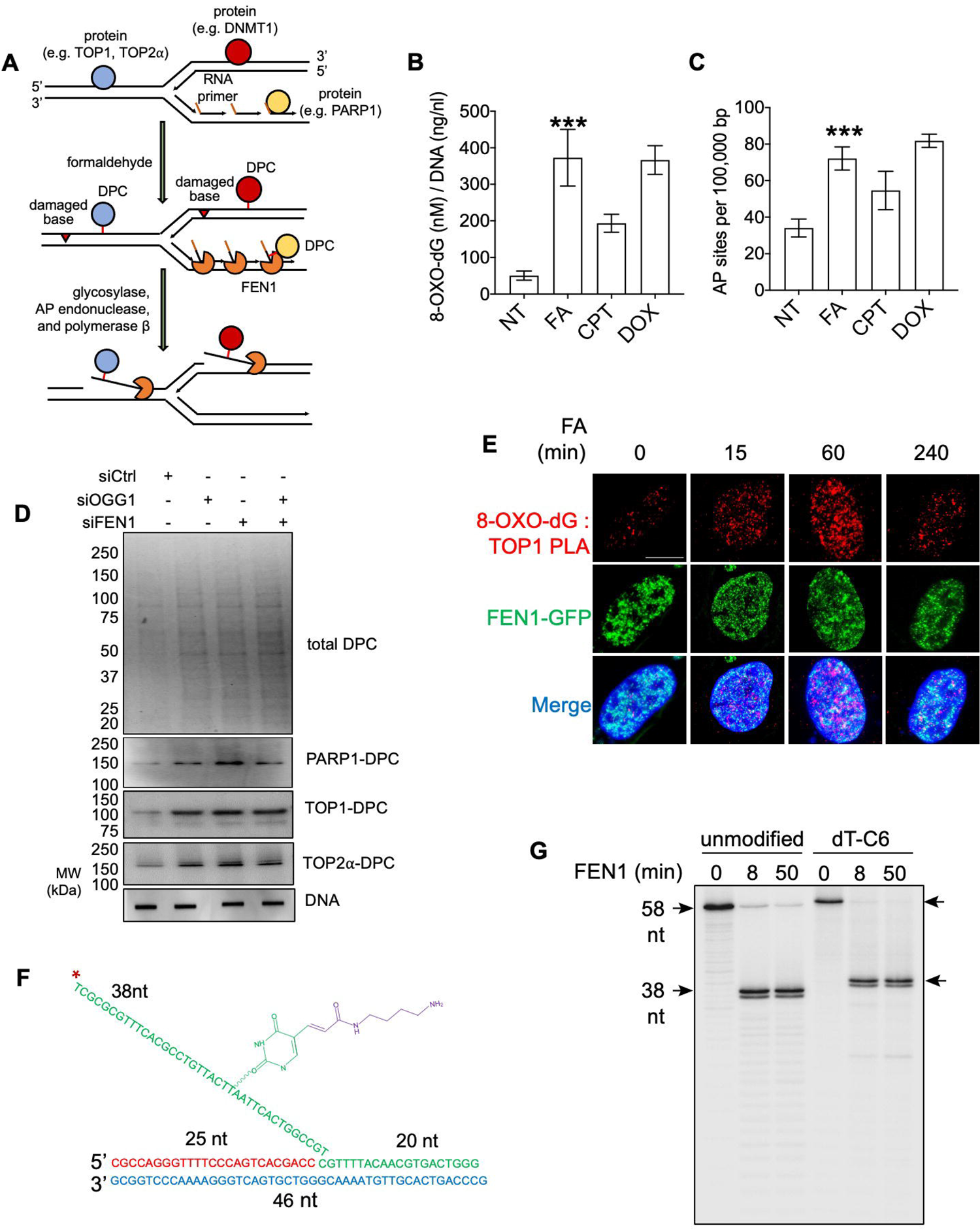
FEN1 cleaves DPC-harboring DNA flap substrate originated oxidative lesions. **A.** Hypothetical model depicting the roles of FEN1 for the repair of FA-induced DPCs in replicating DNA. FA non-specifically crosslinks chromatin-bound proteins in front of the forks and on the leading strand behind the forks as well as proteins that bind Okazaki fragments in lagging strand DNA. DPCs formed within the 5’-flap of Okazaki fragments during strand displacement are readily repaired by FEN1. Concurrent damaged bases adjacent to non-end DPCs (in the front of and on the leading strand behind the replication fork) are converted into DPC-harboring 5’-flaps by the BER proteins, a potential substrate of FEN1. **B.** DNA samples were isolated by RADAR assay in HEK293 cells treated with indicated agents (FA: 400 µM, TOP1 inhibitor campothecin/CPT: 10 µM, TOP2 inhibitor doxorubicin/DOX: 10 µM) for 1 hour. Detection and quantitation of 8-OXO-2’-deoxyguanosine (8-OXO-dG) in the samples were performed using HT 8-oxo-dG ELISA Kit. Error bars, standard deviation (SD). ***, p < 0.001. **C.** Same DNA samples as those in panel c were detected for apurinic or apyrimidinic (AP or abasic) sites using AP Sites Quantitation Kit. **D.** The modified RADAR assay was performed in HEK293 cells transfected with indicated siRNAs and subsequently treated with 400 µM FA for 2 hours. Total DPCs were detected with Coomassie stain and individual DPCs were probed with antibodies targeting the indicated proteins. **E.** Proximity ligation assay (PLA) was performed in FEN1-GFP expression plasmid transfected U2OS cells, following 400 µM FA treatment for indicated periods of time to measure TOP1 and 8-OXO-dG using their respective antibodies. Scale bar represents 10Lμm. **F.** Sequence and structure of double-stranded DNA substrate carrying C6 amino modifier conjugated-5’-flap. *, ^32^P label. **G.** Activity assay testing recombinant human FEN1 towards unmodified and C6-modified DNA substrate shown in panel e for indicated periods of time. ^32^P labeled DNA products following the activity assay were visualized by PAGE electrophoresis.

Prompted by these findings, we next sought to test whether FEN1 can process DNA substrates with 5’-flap-containing DPCs. We designed a DNA substrate with a single-stranded flap internally conjugated with the amino modifier C6 and radiolabeled the 5’-end of the flap with phosphorus-32 (^32^P) to test FEN1 activity towards this substrate (Fig. 3F). Following incubation of recombination FEN1 with unmodified and the amino acid mimetic-conjugated DNA substrates, we ran the samples on a denaturing PAGE and found that FEN1 processed both substrates with comparable efficiency (Fig. 3G). This observation raises the possibility that FEN1 recognizes substrates and directly binds the site of cleavage without tracking along the flap to the junction, which would be interfered by any chemical modifications of individual nucleotides on the flap.

We, therefore, conclude that a 5’-flap DNA strand with an internal nucleotide crosslinked to a peptide can be processed by FEN1 as long as the peptide is away from the point of annealing and does not sterically block the cleavage junction.

### 4. FEN1 prevents FA-induced replication stress and chromosomal instability

Formaldehyde has been implicated in DNA replicative stress and genome instability ^42^. In the absence of FEN1, damaged bases adjoining DPCs are converted into 5’-flaps presumably by the BER pathway but left unrepaired. Upon their collision with ongoing replication forks, single-ended double-strand breaks (seDSBs) are formed and block the replication (Fig. 4A). To determine whether FEN1 relieves FA-induced replication stress, we first measured nascent DNA synthesis by DNA combing assay. Following 1 h incorporation of CIdU into U2OS cells in the presence or absence of FEN1 inhibitor FEN1i, ldU without or without FA or FEN1 inhibitor was added to the cells for 4 h before combing analysis. FEN1 catalytic inhibition alone was found not to perturb fork progression, as supported by our measurement that both CIdU and IdU track lengths were not affected by FEN1 inhibition. FA treatment reduced ldU track length and FA in combination with FEN1 inhibition resulted in a further shortening of the CldU tracks (Fig. 4B; Supplementary fig. 4A).

**Figure 4.**
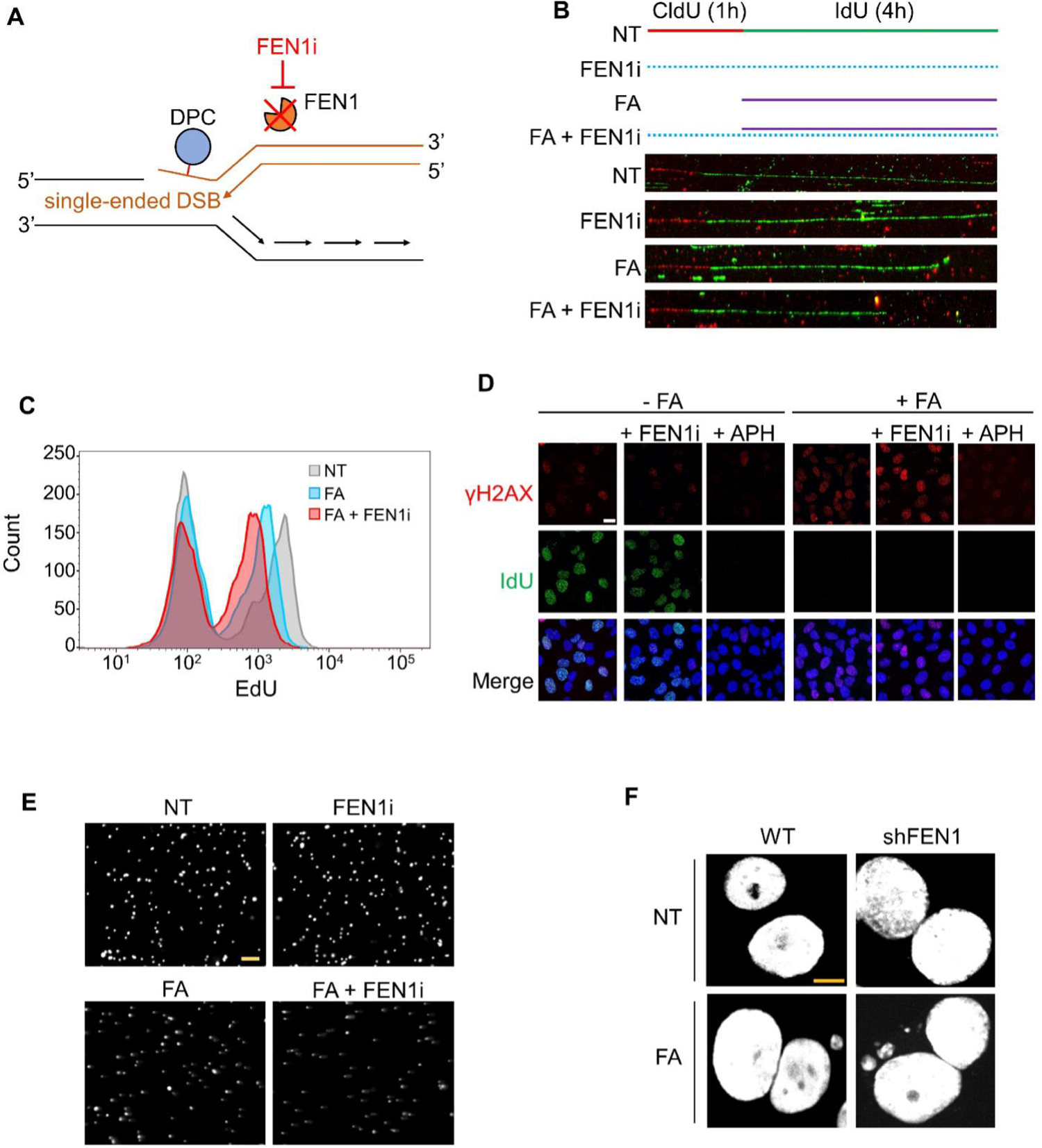
FEN1 prevents FA-induced replication stress and chromosomal instability. **A.** Hypothetical model depicting the induction of DSBs by FA. Ongoing replication forks, once colliding with 5’-flap DPCs ahead of the forks in the absence of FEN1 activity, can be converted into single-ended DSBs that stall replication. **B.** Upper panel: labeling protocols for DNA combing assay in U2OS cells. FEN1i: 10 µM; FA: 400 µM. Lower panel: Representative images of CldU and IdU tracks from combing assays performed under conditions described in the upper panel. **C.** EdU incorporation analyzed by flow cytometry. Cells were treated with FA (400 µM) with or without FEN1i (10 µM) for 4 h and pulsed with EdU (10 µM) for 30 min prior to harvesting. **D.** Representative images of IF of γH2AX and IdU foci by iSIM. U2OS cells were synchronized in the S phase by double thymidine block, followed by IdU incorporation and indicated treatments (400LµM FA, 10 µM FEN1i, 1 µM APH) for 4 hours for IF using anti-γH2AX and anti-BrdU antibodies. **E.** Representative images of neutral comet assay in U2OS cells treated with FEN1i (10LµM, 1Lh), FA (400LµM, 4Lh), and FAL+LFEN1i (pre-treatment with FEN1i for 1Lh then co-treatment for 4Lh). Cells were subjected to neutral comet assay for detection of DNA breaks. Scale bar represents 100Lμm. **F.** Representative images of WT and sh*FEN1* interphase cells treated with or without FA. WT and sh*FEN1* MCF-7 cells were exposed to 500 μM FA for 2 h at 37°C, followed by paraformaldehyde fixation and DAPI staining for microscopic analysis of micronuclei.

To confirm the impact of FA and FEN1 on DNA replication, we performed fluorescence-activated cell sorting (FACS) to quantify the incorporation of EdU during replication. Cells were pre-treated with the FEN1 inhibitor, followed by co-treatment with FA for 4 h and pulse-incorporation of EdU for 30 min. Consistent with the DNA combing assays, FEN1 inhibition alone did not impact EdU uptake (Supplementary fig. 4B). By contrast, FA alone led to a reduction in EdU incorporation, indicative of stalled replication. In line with the combing assay, FA combined with FEN1 inhibition further decreased EdU incorporation (Fig. 4C). Together, these results demonstrate that FEN1 is a critical factor limiting FA-induced replication stress presumably by removing the DPCs generated by FA.

Persistent DPCs formed ahead of replication forks physically obstruct replication fork progression, leading to fork collapse, DSBs, and ultimately chromosomal instability. To test the role of FEN1 in limiting FA-induced DNA damage, we measured γH2AX foci by immunofluorescence (IF) microscopy and found that FA treatment induced γH2AX formation and blocked IdU incorporation and that FEN1 inhibition enhanced FA-induced γH2AX accrual (Fig. 4D; Supplementary fig. 4C). Replication inhibition by aphidicolin (APH) inhibited γH2AX induction by FA, indicating the requirement of replication for the induction of DNA damage by FA. In parallel, we conducted neutral comet assays in U2OS cells and found that FEN1 inhibition exacerbated FA-induced DNA single- and double-strand breaks (Fig. 4E; Supplementary fig. 4D). To determine whether FEN1 plays a role in FA-induced chromosomal instability, we assessed micronuclei that arise from lagging acentric chromosome or chromatid fragments in anaphase by microscopy and found a significant elevation in micronucleus levels in FEN1 KD MCF-7 cells (Fig. 4F; Supplementary fig. 4E). Together, these data demonstrate a key role of FEN1 in preventing FA-induced replication fork stalling, replicative DNA damage and chromosomal instability likely by efficiently repairing FA-induced DPCs.

### 5. FEN1 repairs enzymatic TOP2-DPCs independently of the BER pathway

To determine if FEN1 also repairs enzymatic DPCs, we performed the ICE assay in FEN1-deficient human cells treated with TOP1 inhibitor camptothecin (CPT) and TOP2 etoposide (ETOP) to induce enzymatic TOP1 and TOP2-DPCs, two of the most studied enzymatic DPCs, respectively. In line with that TOP1 acts by generating 3’ DNA cleavage complexes whereas TOP2 forms 5’ DNA cleavage complexes, it was found that FEN1 downregulation accumulated ETOP-induced enzymatic TOP2 (both α and β isozymes) - DPCs but not CPT-induced enzymatic DPCs (Fig. 5A and B; Supplementary fig. 5A). This observation is consistent with a previous finding that FEN1 may possess nucleolytic activity against single-stranded 5’ phoshotyrosyl termini^43^.

**Figure 5.**
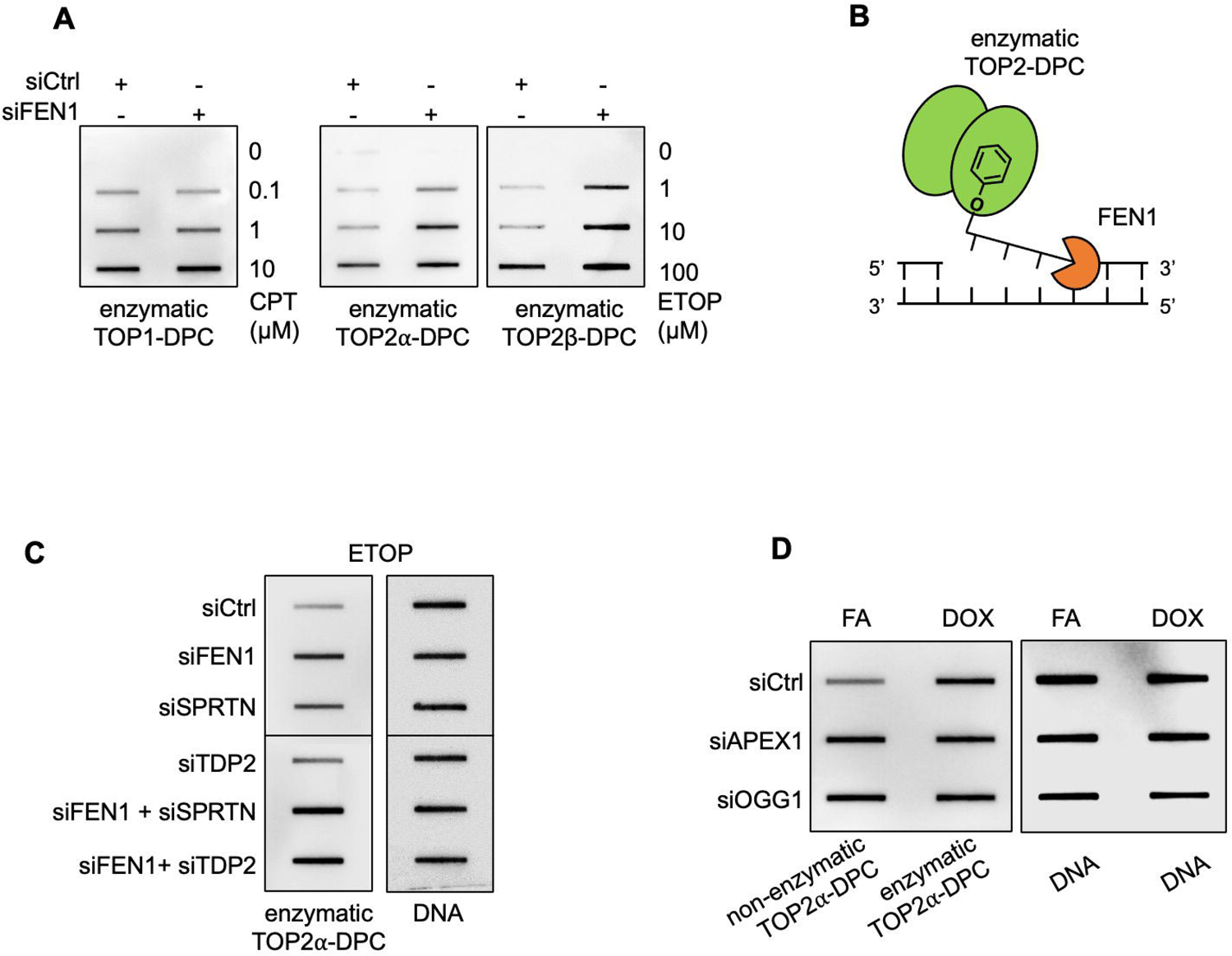
FEN1 repairs enzymatic TOP2-DPCs independently of the BER pathway. **A.** The ICE assay in U2OS cells treated with indicated agents (camptotehcin, CPT; etoposide, ETOP) for 30 min before immunostaining with indicated antibodies. **B.** Hypothetical model depicting the repair of enzymatic TOP2-DPCs by FEN1. The single-stranded 5’ TOP2-DPC induced by TOP2 inhibitors such as etoposide or doxorubicin (DOX) can be converted to a 5’ flap structure via post-translational modifications, conformational changes, or polymerase activities for FEN1 to process. **C.** The RADAR assay in HEK293 cells transfected with indicated siRNAs for 48 h then treated with ETOP at 10 µM for 1 h. Enzymatic TOP2α-DPCs were probed with TOP2α antibody and DNA was probed with anti-DNA antibody. **D.** The RADAR assay in HEK293 cells transfected with indicated siRNAs for 48 h then treated with FA at 1 mM or DOX at 10 µM for 1 h. TOP2α-DPCs were probed with TOP2α antibody and DNA was probed with anti-DNA.

Double knocking-down of FEN1 and SPRTN and double knocking-down of FEN1 and TDP2, a tyrosine 5-phosphodiesterase, both stimulated ETOP-induced enzymatic TOP2-DPCs in comparison with their respective single knocking-down (Fig. 5C; Supplementary fig. 5B), suggesting FEN1 as a salvage pathway in parallel to SPRTN and TDP2. As FEN1-dependent repair of FA-induced non-enzymatic DPCs requires the BER pathway to convert the no-break DPCs into 5’ flap DPCs, we next investigate whether FEN1-mediated enzymatic TO2-DPC repair also involves the BER pathway. As expected, the RADAR assay found downregulation of APE1 or OGG1 elevated FA-induced nonenzymatic TOP2-DPC levels (Fig. 5D; Supplementary fig. 5C). Deficiency in APE1 or OGG1, on the contrary, did not affect enzymatic TOP2-DPCs induced by doxorubicin, a TOP2 inhibitor that also induces oxidative damage (see Fig. 3B and C). This finding suggests that FEN1 acts on enzymatic TOP2-DPCs with 5’ broken ends without the need for the BER pathway.

### 6. PARP1 regulates FEN1-dependent repair of non-enzymatic and enzymatic DPCs

After having established the role of FEN1 in the repair of non-enzymatic DPCs and the prevention of DPC-induced genotoxic events, we sought to identify the molecular mechanisms regulating FEN1 in DPC repair. As our ICE-MS profiling identified PARP1 DPCs among the most abundant DPCs induced by FA and our RNAi screen discovered PARP1 as a resistance gene to FA cytotoxicity, we examined the role of PARP1 and PARylation in regulating FEN1-meidated DPC repair. PARP1 trapping or binding to damaged DNA elicits *in trans* autoPARylaion and PARylation of the damaged sites/chromatin and DNA repair proteins, which collectively promote the repair of DNA lesions including single- and double-strand breaks by homologous recombination or non-homologous end-joining, nucleotide excision repair (NER), as well as BER ^37, 44^.

We hypothesized that FA induction of PARP1-DPCs as well as other nonenzymatic DPCs activates PARP1 and that PARylation signals repair proteins to be recruited to the DPC sites. To test this hypothesis, we first tested whether FA-induced DPCs are PARylated. Using the modified RADAR assays in HEK293 cells treated with FA along with several other DPC inducers, we failed to detect any PARylation signal on these DPCs. However, pre-treatment with PARG (PAR glycohydrolase) inhibitor ^45–47^ led to readily detectable PARylation of the DPCs induced by FA but not of those induced by the other DPC inducers tested (Fig. 6A; Supplementary fig. 6A). We also observed upshifting of the DPC signal (indicative of PARylated DPC of higher molecular weight and hence lower electrophoretic mobility) (Fig. 6A). These results demonstrate that FA-induced DPCs are specifically and reversibly PARylated. Akin to FA-induced non-enzymatic DPCs, enzymatic TOP2-DPCs induced by ETOP appeared to be PARylated and promptly revered by PARG, as evidenced by our observation that TOP2-DPC PARylation became detectable when PARG was blocked (Fig. 6B).

**Figure 6.**
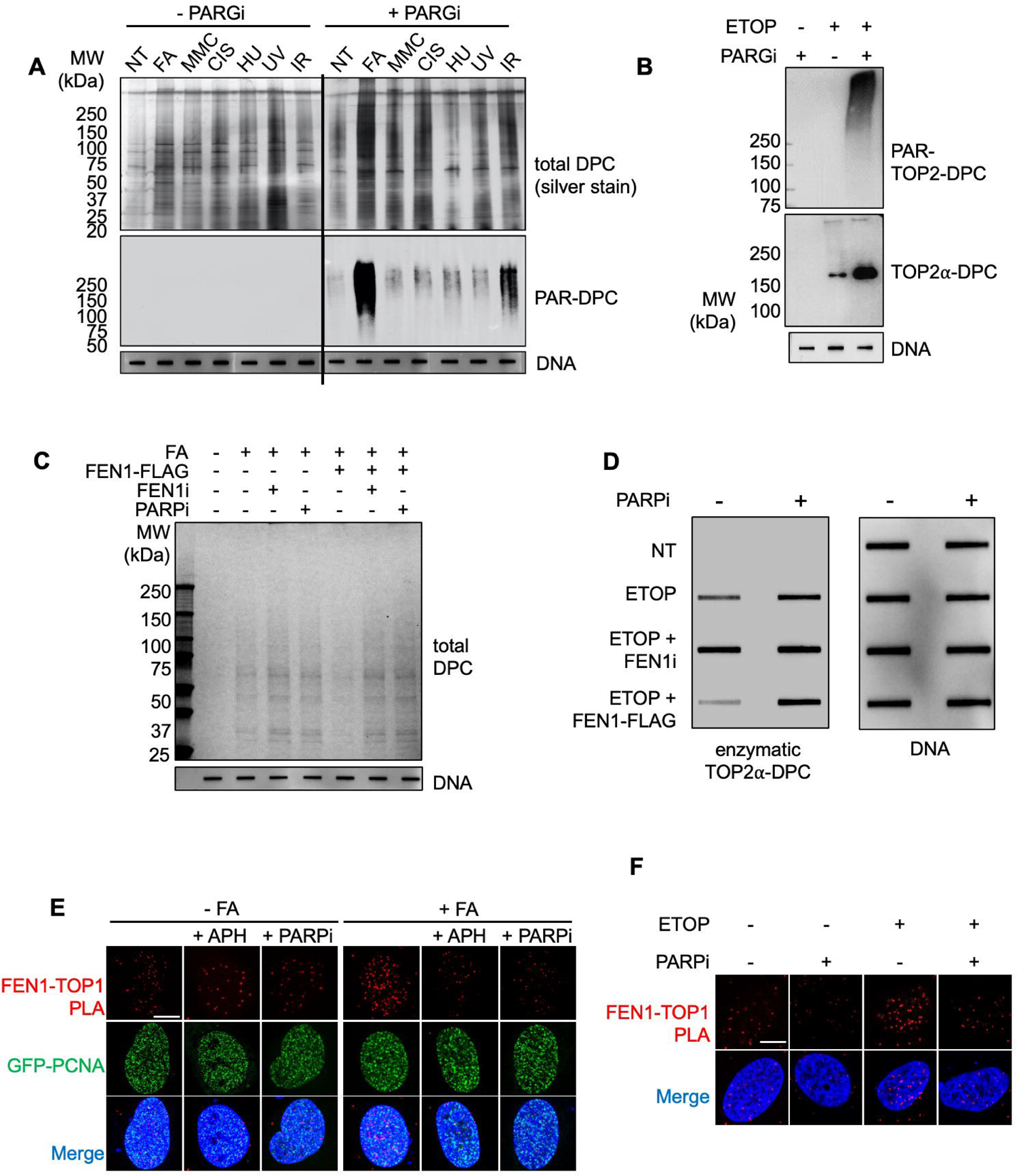
PARP1 regulates FEN1-dependent repair of non-enzymatic and enzymatic DPCs. **A.** The modified RADAR assay in HEK293 cells treated with indicated agents. FA: 400 µM for 2 h; HU: 1 mM for 2 h; MMC: 10 µM for 2h; Cisplatin: 10 µM for 2 h; UV: 20 J/m2; IR: 20 Gy. Cells were exposed to 10 µM PARGi for 1h before co-treatments. Total DPCs were detected by silver stain and PARylated DPCs were probed with anti-PAR antibody. **B.** The modified RADAR assay in HEK293 cells treated with indicated agents. PARGi was used at 10 µM for 1 h pre-treatment then 1 h co-treatment with 10 µM ETOP. **C.** HEK293 cells were transfected with empty vector or FEN1-FLAG overexpression plasmid, followed by treatments with indicated agents. FA: 400 µM for 2 h; FEN1i: 10 µM, 1 h pre-treatment + 2 h co-treatment with FA; PARPi: 10 µM, 1 h pre-treatment + 2 h co-treatment with FA. Cells were subjected to the modified RADAR assay for detection of total DPCs by Coomassie stain. **D.** The RADAR assay in HEK293 cells transfected with or without FEN1-FLAG expression plasmid for 48 h then treated with indicated agents. PARPi and FEN1i were used at 10 µM for 1 h pre-treatment, followed by co-treatment with ETOP at 10 µM for 1 h. Enzymatic TOP2α-DPCs were probed with TOP2α antibody and DNA was probed with anti-DNA antibody. **E.** GFP-PCNA expression plasmid transfected U2OS cells were treated with indicated agents (FA: 400 µM for 1 h; APH: 1 µM, 30 min pre-treatment + 1 h co-treatment with FA; PARPi: 10 µM, 1 h pre-treatment + 1 h co-treatment with FA). Cells were then subjected to PLA assay to measure TOP1 and FEN1 interaction using their respective antibodies. Scale bar represents 10Lμm. **F.** U2OS cells were treated with indicated agents (ETOP: 10 µM for 1 h; PARPi: 10 µM, 1 h pre-treatment + 1 h co-treatment with ETOP). Cells were subjected to PLA assay to measure TOP2α and FEN1 interaction using their antibodies. Scale bar represents 10Lμm.

Next, we assessed whether FEN1 can process PARylated DPCs by incubating RADAR samples from cells exposed to FA in the presence of PARG inhibitor with recombinant FEN1 and found that FEN1 released PARylated DPCs induced by FA plus PARG inhibitor (Supplementary fig. 6B). This result shows that PARylation does not interfere with FEN1 activity and led us to hypothesize that PARylation of the DPCs serves as a recruitment signal, as PAR-branched polymers resemble nucleic acid molecules.

To further investigate the potential role of PARP1 in the repair of non-enzymatic DPCs, we performed the modified RADAR assay in HEK293 cells transfected with FEN1 overexpression plasmid and observed that upregulation of FEN1 facilitated the removal of FA-induced DPCs whereas the facilitation was reversed by FEN1 inhibition and by PARP inhibition by talazoparib (PARPi) (Fig. 6C; Supplementary fig. 6C). It was also found by the RADAR assay that enzymatic TOP2-DPCs induced by ETOP were cleared by upregulation of FEN1, and that the FEN1-dependent removal was inhibited by talazoparib (Fig. 6D; Supplementary fig. 6D). Further, PLA assays showed that PARP1 inhibition diminished FA-induced TOP1 and FEN1 interaction as did aphidicolin (Fig. 6E; Supplementary fig. 6E). Similarly, ETOP-induced TOP2α-FEN1 colocalization was also disrupted by PARP1 inhibition (Fig. 6F; Supplementary fig. 6F). Taken together, these data demonstrate the role of PARP1 and PARP1 PARylation in FEN1-mediated DPC repair.

### 7. ADP-ribosylation of FEN1 at glutamic acid residue 285 recruit FEN1 to DPC sites

To determine whether FEN1 is a direct substrate of PARP1, we conducted PAR antibody-based nuclear immunoprecipitation (IP) under denaturing conditions in HEK293 cells treated with PARG inhibitor for mass spectrometry analysis (Fig. 7A; Supplementary fig. 7A). We identified FEN1 as a PARylation substrate and confirmed the result by *in vitro* PARylation assay using recombinant FEN1 and PARP1 proteins (Fig. 7B). Further, FLAG-IP in HEK293 cells transfected with FLAG-FEN1 stimulated FEN1 PARylation by FA, and PARG inhibition further enhanced FA-stimulated FEN1 PARylation (Supplementary fig. 7B).

**Figure 7.**
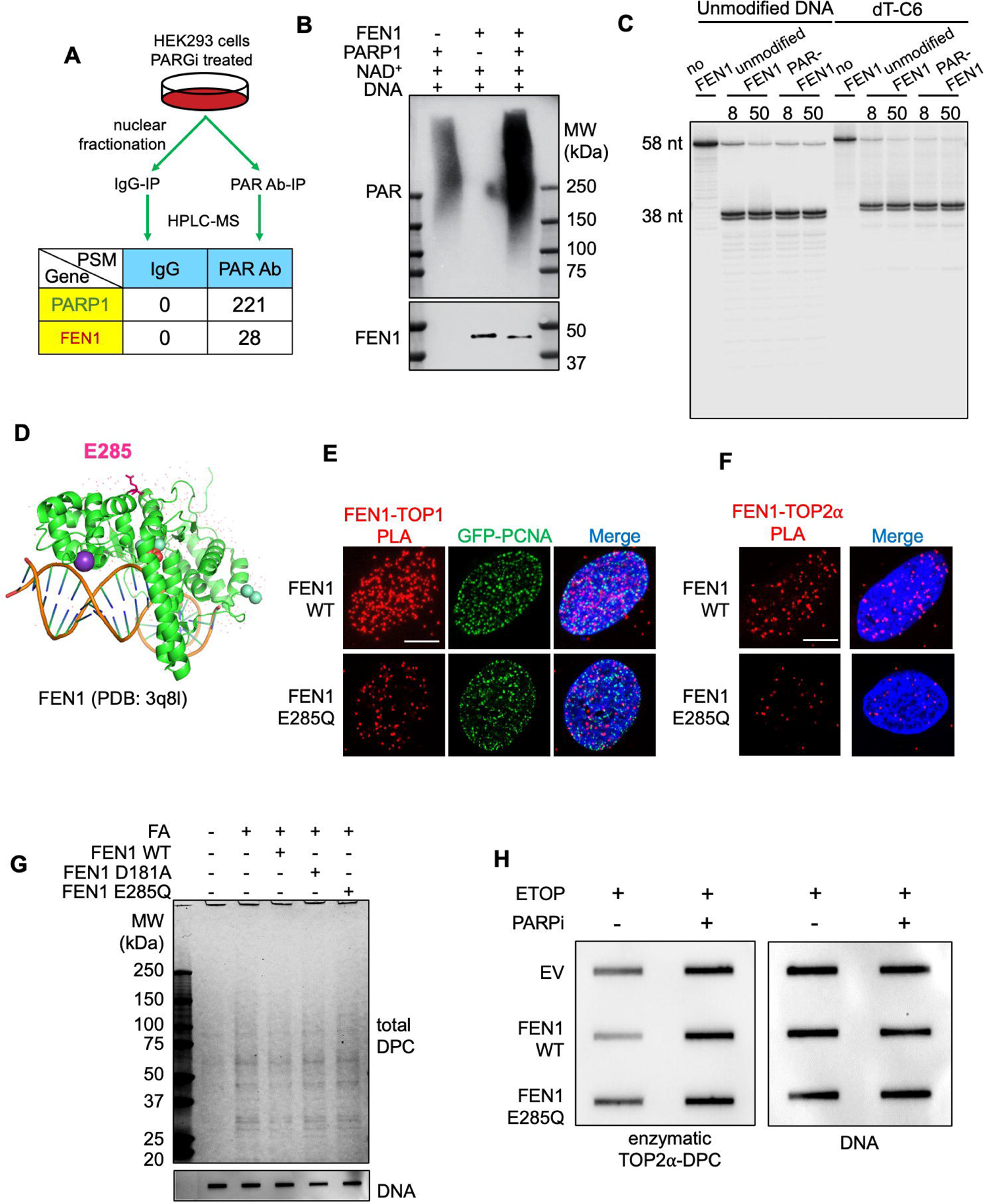
E285 ADP-ribosylation localizes FEN1 to DPC sites for repair. **A.** Experimental setup for PAR antibody-IP-MS in HEK293 cells. PSM, peptide spectrum match. **B.** *in vitro* PARylation assay with recombinant FEN1 protein, recombinant PARP1 protein, NAD^+^ and activated DNA. Following 20 min incubation at room temperature, samples were subjected to Western blotting using anti-PAR and anti-FEN1 antibodies. anti-FEN1 antibody failed to detect PARylated FEN1 likely because PAR polymers blocked the epitope. **C.** Activity assay testing unmodified and PARylated recombinant FEN1 proteins towards unmodified and C6-modified DNA substrate indicated periods of time. ^32^P labeled DNA products following the activity assay were visualized by PAGE electrophoresis. **D.** Structure of FEN1 with DNA substrate, SM^3+^ and K^+^ (PDB: 3q8l) highlighting glutamic acid residue 285 (E285), an ADP-ribosylation site identified by ELTA-MS. **E.** GFP-PCNA expressing U2OS cells transfected with FEN1-FLAG WT or FEN1-FLAG E285Q expression plasmid were treated with 400 µM FA for 1 h. Cells were then subjected to PLA assay to measure TOP1 and FEN1-FLAG interaction using their TOP1 and FLAG antibodies. Scale bar represents 10Lμm. **F.** U2OS cells transfected with FEN1-FLAG WT or FEN1-FLAG E285Q expression plasmid were treated with 10 µM ETOP for 1 h. Cells were then subjected to PLA assay to measure TOP2α and FEN1-FLAG interaction using TOP2α and FLAG antibodies. Scale bar represents 10Lμm. **G.** HEK293 cells were transfected with empty vector or indicated FEN1-FLAG overexpression plasmid, followed by treatment with 400 µM FA for 2 h. Cells were subjected to the modified RADAR assay for detection of total DPCs by Coomassie stain. **H.** HEK293 cells were transfected with empty vector or indicated FEN1-FLAG overexpression plasmid, followed by treatment with 10 µM ETOP for 30 min. Cells were subjected to the RADAR assay for detection of enzymatic TOP2α-DPCs and DNA using their antibodies.

We next performed *in vitro* PARylation of recombinant FEN1 and tested the modified FEN1 in activity assay using the 5’-flap DNA substrate (see Fig. 3F). Unmodified and PARylated FEN1 exhibited comparable efficiency against the DNA substrates (both unmodified and amino modifier C6-conjugated) (Fig. 7C). This observation suggests that PARylation of FEN1 plays a role in recruiting FEN1 to the DPC sites without affecting the catalytic activity of FEN1 towards to 5’-flap DPCs.

To test the hypothesis that FEN1 PARylation by PARP1 drives FEN1 to DPCs, we applied the ELTA (Enzymatic Labeling of Terminal ADP-Ribose) method for mass spectrometry identification of the ADP-ribosylation sites on FEN1 ^48^. In brief, we first PARylated recombinant FEN1 *in vitro* using PARP1 and NAD^+^. Following trypsin digestion, we labeled FEN1 peptide-conjugated ADP-ribose polymers at their 2’-OH termini using the enzyme OAS1 and dATP. After enrichment of the labeled FEN1 peptide-ADP-ribose conjugates, we treated the sample with NudT16, which cleaves the pyrophosphate bond within ADP-ribose to leave a phosphoribosyl group on protein residues that were previously ADP-ribosylated, for MS profiling of the ADP-ribosylation sites. With this method we identified glutamic acid residue 285 (E285) as a major ADP-ribosylation site of FEN1 (Fig. 7D). We next mutated E285 to glutamine (E285Q) to characterize this modification site using the PLA assay and found that the mutating E285 led to a significant reduction in FA-induced TOP1-FEN1 interaction foci and ETOP-induced TOP2α-FEN1 interaction foci (Fig. 7E and F; Supplementary fig. 7C and D). This result with E285 PARylation acting as a recruitment mechanism for the localization of FEN1 to DPCs. In consistence with observation, the modified RADAR assay showed that upregulation FEN1 E285Q or the nuclease-dead mutant D181A failed to promote the removal of FA-induced DPC, as opposed to the native FEN1 counterpart (Fig. 7G; Supplementary fig. 7E). In consistence, FEN1 E285Q was also found by the RADAR assay unable to remove ETOP-induced enzymatic TOP2α-DPCs (Fig. 7H; Supplementary fig. 7F). In sum, these findings point to a model in which PARP1 is activated upon the DPC-conjugated 5’-flap induction and modifies FEN1 with PAR to mediate FEN1 localization to the DPCs.

## Discussion

In the present study, we systematically profiled proteins crosslinked to DNA by FA using ICE-MS and identify FEN1 as a previously unrecognized nuclease excising FA-induced DPCs downstream from a PARylation pathway. FA non-specifically crosslinks chromatin-bound proteins to DNA bases by forming a secondary amine (-NH-CH-NH-) bonding the proteins to the DNA backbone. Because the crosslinking is refractory to peptidases that only digest amide (peptide) bonds, we speculated that the cell must engage a nuclease(s) to remove the full-length DPCs or the residual DNA-peptide crosslinks resulting from partial proteolysis. As opposed to enzymatic DPCs such as the topoisomerase DPCs that are linked to one (or two) end of the broken nucleic acid, FA crosslinks proteins to DNA without introducing terminal breaks in the DNA, raising the possibility that repair nuclease(s) must incise the DNA backbone adjacent to the DPCs.

FEN1 plays crucial roles in DNA replication and repair by processing 5’-flap structures that arise from strand displacement synthesis by Pol δ during Okazaki fragment maturation and from displacement of DNA broken ends by Pol δ, Pol ε or Pol β during long-patch BER pathway ^28, 49^. Based on the roles of FEN1 in DNA replication and repair, we propose that FEN1 processes DPCs formed ahead of the replication forks, on the leading strand (or its parental stand) behind the forks, as well as on the flap of Okazaki fragments (lagging strand) (Fig. 8A). In order for FEN1 to cleave DPCs in front of the fork, it can be postulated that 5’-flap structures are likely formed as a byproduct of BER processing of damaged bases induced in close proximity to the DPCs. Indeed, we observed induction of base damage by FA co-localizing with DPCs formed ahead of replication forks (e.g. non-enzymatic TOP1-DPCs), supporting our above-mentioned model for FEN1-mediated repair of DPCs formed ahead of the replication fork (Fig. 8A). As a result, FEN1 was found to attenuate FA-induced replication stress to ensure DNA synthesis with its endonuclease activity.

Importantly, we show in the study that FEN1 repairs enzymatic TOP2-DPCs independently of the BER pathway (Fig. 8B), as opposed to its role in the repair of FA-induced non-enzymatic DPCs. Although TOP2 homodimers cleave both strands of the DNA duplex to generate the enzyme-DNA covalent intermediates (TOP2 cleavage complexes or TOP2ccs) and religate the broken end after strand passage, religation of the two strands are independently inhibited by TOP2 inhibitors ^50^. It has been reported a large fraction of DNA breaks induced by TOP2 inhibitors including ETOP appears to be protein-linked single-strand breaks (SSBs) ^51–53^. These 5’ single-strand DPCs can be converted into 5’ flap structures presumably through conformational alteration in the protein or polymerase activity without the BER pathway as a precondition for its cleavage.

One earlier study shows that perturbation of replication by downregulating FEN1 enhances PARP activity^46^. Yet, our study demonstrates PARP1-mediated ADP-ribosylation as a key mechanism modulating the recruitment of FEN1 to DPCs. We discovered that both DPCs (FA-induced non-enzymatic DPCs and ETOP-induced enzymatic TOP2-DPCs) and FEN1 are PARylation substrates and that PARylation is promptly reversed by PARG and hence highly transient. We propose that DPC PARylation by PARP1 acts as a signal to recruit FEN1 with PARylation acting as an interaction scaffold. The likely subsequent PARylation of FEN1 recruits it to the PARylated DPCs, followed by PARG-dependent dePARylation of both the DPCs and FEN1 to enable FEN1-catalyzed cleavage of the DPCs (Fig. 8A and B). Such a model is supported by our finding that FEN1 E285 PARylation is required for its co-localization with fork-front DPCs such as TOP1 and TOP2α ^7, 8^. Given that FEN1 is enriched near Okazaki fragments for their maturation, and that DPCs formed within the flap of Okazaki fragments can be easily detected by PARP1 and cleaved by FEN1, it is plausible that PARylation drives the translocation of FEN1 from the rear of the fork to its front or to the leading strand behind the fork to repair DPCs formed within those two regions (Fig. 8A).

It has been implicated that proteolytic digestion of enzymatic DPCs (e.g. TOP-DPCs) enables or facilitates the processing of the phosphotyrosyl bond between the enzyme and DNA by tyrosyl-DNA phosphodiesterases ^19, 20^. This is the case for FEN1-mediated repair of non-enzymatic DPCs and enzymatic TOP2-DPCs, as demonstrated by our observation that downregulation of FEN1 and SPRTN markedly accumulated non-enzymatic DPCs and enzymatic TOP2-DPCs in comparison with their respective single downregulation. Also, FEN1 single knocking-down was found to elevate full-length non-enzymatic DPCs. If partial proteolysis of the DPCs by either SPRTN or proteasome was a precondition for FEN1-dependent cleavage of DPCs, partially proteolyzed DPCs instead of the full-length counterpart should have accumulated upon FEN1 single knocking-down. Our findings directly suggest that FEN1 and the SPRTN protease serve as parallel pathways for both non-enzymatic and enzymatic DPC repair. The partially proteolyzed enzymatic TOP2 DPCs are repaired by TDP2, the 5’ tyrosine-DNA phosphodiesterase, whereas FA-induced DPCs might be repaired by other nucleolytic mechanisms or by homologous recombination during replication following their proteolysis. Yet, how transcription-associated DPCs are repaired remains to be clarified. Our preliminary finding that downregulation of XPF (ERCC4) or XPG (ERCC5) conferred hypersensitivity to FA and accumulated FA-induced DPCs in the cell suggests an involvement of transcription-coupled nucleolytic excision repair (TC-NER) in the repair of general non-enzymatic DPCs and hence warrants further investigation.

In conclusion, our work discovered an unprecedented nucleolytic pathway as a universal repair mechanism for non-enzymatic and enzymatic DPCs from different sources and a central factor for the avoidance of DPC-induced genome instability. Defects in (both enzymatic and non-enzymatic) DPC repair have been implicated as a major cause of progeroid features, cancers including hepatocellular carcinoma, neurodegeneration as well as immunodeficiencies ^6, 19^. Consistently, somatic FEN1 mutations can lead to autoimmunity, chronic inflammation, and cancers ^28, 54^. Our work provides previously unrecognized pieces to the puzzle of DPC repair, and molecular foundation for the etiology of DPC-induced diseases.

## Methods and Materials

### Human Cell Culture

Human embryonic kidney HEK293 cells, human breast cancer MCF7 cells and human bone osteosarcoma U2OS cells were in cultured in DMEM medium (Life Technologies) supplemented with 10% (v/v) fetal bovine serum, 100 units/ml penicillin, 100 μg streptomycin /ml streptomycin and 1x GlutaMax in tissue culture dishes at 37 °C in a humidified CO2 – regulated (5%) incubator.

### Chemicals

Formaldehyde solution (FA, Sigma Aldrich); FEN1-IN-4 (FEN1i, Selleck); PDD 00017273 (PARGi, Tocris); the replication inhibitor aphidicolin (APD, Sigma Aldrich); Etoposide (ETOP, Sigma Aldrich); talazoparib (PARPi, Selleck). AcquaStain protein gel Coomassie stain (Bulldog Bio); Silver Stain solutions (Bio-Rad).

### Antibodies

#### The following antibodies were used

anti-PAR, mouse monoclonal, Trevigen, 4335-MC-100; anti-PAR (10H), mouse monoclonal, Enzo Life Sciences, ALX-804-220; anti-TOP1, mouse monoclonal, BD Biosciences, 556597; anti-dsDNA, mouse monoclonal, Abcam, ab27156; anti-FLAG, mouse monoclonal, Sigma Aldrich, F1804; anti-FLAG, rabbit polyclonal, Sigma Aldrich, F7425; anti-FEN1, rabbit polyclonal, Cell Signaling, 2746; anti-XPF, rabbit monoclonal, 13465; anti-XPG, mouse monoclonal, Santa Cruz, 13563; anti-APE1, rabbit monoclonal, Cell Signaling, 10519; anti-EXO1, rabbit polyclonal, Abcam, 95068, anti-DNA2, rabbit polyclonal, Abcam, 96488; anti-MRE11, mouse monoclonal, GeneTex, 70212; anti-TDP1, rabbit polyclonal, Bethyl Laboratories, A301-618A; anti-TDP2, mouse monoclonal, Santa Cruz, 377280; anti-PARP1 (F2), mouse monoclonal, Santa Cruz, sc-8007; anti-PARG, rabbit monoclonal, Cell Signaling, 66564; anti-γH2AX, rabbit polyclonal, Cell Signaling, 66564; anti-BrdU, mouse monoclonal, ab8152.

#### Recombinant proteins

Human recombinant FEN1 is a gift from Dr. Rajendra Prasad at NIEHS. Human recombinant PARP1 was purified from *E. coli* as described ^55^. Human 2’−5’-oligoadenylate synthase 1 (OAS1) was purified from SF9 insect cells as described.

### Expression plasmids

**The following expression plasmids were used:** pShuttle-FLAG-FEN1, Addgene, 35027; pCMV-6×His-TOP1-HaloTag ^17^; pCMV PARP1-3×Flag, Addgene, 111575; pCMV6-AC-FEN1-turboGFP, OriGene, RG201785; pCMV6-AC-PCNA-turboGFP, OriGene, RG201741. 48 h transfection was performed using lipofectamine™ 3000 transfection reagent (Invitrogen) following the manufacturer’s instructions.

### Small-interfering RNA (siRNA) and small-hairpin RNA (shRNA)

**The following expression plasmids were used:** control siRNA, Dharmacon, D-001206-13-05; FEN1 siRNA, Dharmacon, M-010344-01; ERCC4 (XPF) siRNA, Dharmacon, M-019946-00; ERCC5 (XPG) siRNA, Dharmacon, M-006626-01; APEX1 (APE1) siRNA, Dharmacon, M-010237-01; DNA2 siRNA, Dharmacon, M-026431-01; EXO1 siRNA, Dharmacon, M-013120-00; MRE11 siRNA, Dharmacon, M-009271-01; TDP1 siRNA, Dharmacon, M-016112-01; TDP2 siRNA, Dharma, M-017578-00. 72 h transfection was performed using lipofectamine™ RNAiMAX transfection reagent (Invitrogen) following the manufacturer’s instructions.

### Lentivirus-mediated gene silencing of FEN1

shRNA oligonucleotides targeting FEN1(shown in the following table) were cloned into pLKO.1 (Cat# 8453, Addgene) digested with EcoRI and AgeI. pLKO.1-shFEN1 or pLKO.1 control vector was simultaneously transfected into LentiX293T (Cat# 632180 Clontech, Japan) with packaging plasmid, pSPAX2 (Cat# 12260, Addgene) and envelop plasmid, pMD2.G (Cat# 12259, Addgene). After harvesting the medium (3 ml) containing lentiviral particles, we enriched the lentiviral particles with Lent-X Concentrator (Cat# 631231, Clontech, Japan) according to the manufacturer’s protocol. The supernatant-containing virus was mixed with wild-type MCF-7 cells. The infected cells (Puromycin resistant) were enriched by puromycin drug selection for 72 hr. Downregulation of FEN1 expression was confirmed by western blotting using anti-FEN1 antibody.

Oligonucleotides (shRNA sequences for pLKO.1 vector) FEN1, forward primer, 5’-CCGGGATGCCTCTATGAGCATTTATCTCGAGATAAATGCTCATA GAGGCATCTTTTTG-3’ FEN1, reverse primer, 5’-AATTCAAAAAGATGCCTCTATGAGCATTTATCTCGAGATAAAT GCTCATAGAGGCATC-3’

### RNA interference (RNAi) high-throughput screening

A focused screen was performed using the Dharmacon ON-TARGETplus® Human DNA Damage Response siRNA library (Horizon Discovery GU-106005-025) targeting 239 genes with 4 distinct siRNAs/target, 1 siRNA/well for a total of 956 siRNAs arrayed into three 384 well plates. MCF7 cells (750 cells/well) were reverse transfected using Lipofectamine RNAiMAX (0.15ul/well, Thermofisher 13778150) with a final siRNA concentration of 20nM. Two sets of screens were performed in parallel. 24 hours post transfection, one set was treated with 80 LM formaldehyde (FA), the other with vehicle DMSO, for an additional 3 days. On day 4, cell numbers were evaluated using CellTiter-Glo® One luminescence (Promega G8462) read on a BMG Pherastar FSX plate reader.

Screen performance was evaluated using AllStars Hs cell death control siRNA (Qiagen) compared to non-targeting control siRNA#2 (Thermofisher). Screen z’ averaged 0.7, indicating excellent screen performance. Ratios of luminescence of FA-treated wells normalized to median luminescence of FA-treated NT siRNA wells over luminescence of vehicle-treated wells normalized to median luminescence of vehicle-treated NT siRNA wells were calculated to determine the combined effect of siRNA knockdown and FA treatment. These ratios were transformed to z scores using mean (μ) and standard deviation (σ), z = (x-μ)/σ. Genes with negative median z score of the 4 siRNAs were considered to be targets whose knockdown sensitized cells to FA-induced damage.

### Western blotting

Cellular proteins were detected by lysing cells with RIPA buffer (150 mM NaCl, 1% NP-40, 0.5% Sodium deoxycholate, 0.1% SDS, 50 mM Tris pH 7.5, 1 mM DTT and protease inhibitor cocktail), followed by sonication and centrifugation. The supernatant was collected and boiled for 10 mins, analyzed by SDS-PAGE, and immunoblotted with various antibodies as indicated. TOP1 downregulation was monitored using alkaline lysis method as described previously.

### Viability assay

10,000 cells were seeded in 96-well white plates (PerkinElmer Life Sciences, 6007680) in triplicate in 100 μl of medium per well overnight. The next day, cells were exposed to drugs and incubated for 72 hours. Cellular viability was determined using the ATPlite 1-step kits (PerkinElmer). 50 µl of ATPlite solution was added in 96-well plates for 15 min, followed by luminescence measurement with an EnVision 2104 Multilabel Reader (PerkinElmer). The adenosine 5′-triphosphate (ATP) level in untreated cells was defined as 100%. Viability (%) of treated cells was defined as ATP-treated cells/ATP-untreated cells × 100.

### *In vitro* FEN1 PARylation

Recombinant FEN1 (5 μg) was incubated with 200 nM recombinant PARP1 enzyme (5 μg) in 1 × PARylation buffer (50 mM Tris-HCl pH 8.0, 50 mM NaCl, 10 mM MgCl_2_, 2% glycerol, 1 mM DTT). 1 mM NAD^+^ were added to the reaction as indicated. The reactions were incubated at room temperature for 20 min and inactivated by adding talazoparib (1 μM) for ELTA-MS analysis or by adding SDS sample buffer for Western blotting analysis.

### FEN1 activity assay

**The following oligonucleotides were used in the study:** D_amino_ (downstream primer that forms a 38 nt flap) 5’-TCGCGCGTTTCACGCCTGTTACTTAATT*CACTGGCCGTCGTTTTACAACGTGACTGGG* The 5-methyl group in thymine of T28 is modified with the amino modifier C6. U1 (upstream primer) 5’-CGCCAGGGTTTTCCCAGTCACGACC Tamino (template) 5’-GCCCAGTCACGTTGTAAAACGGGTCGTGACTGGGAAAACCCTGGCG

The 3‘-end regions of the downstream primer D_amino_ is homologous with the 5‘-end of its template. Once annealed, these primers create substrates with unannealed 5‘-tails as shown in the figures. The upstream primer U1 was annealed to its template to create a nick at the base of the unannealed 5‘-tail of the downstream primer D_amino_. Prior to annealing, the downstream primer D_amino_ was 5‘-phosphorylated and radiolabeled using [γ-32P]ATP (PerkinElmer) and T4 polynucleotide kinase (NEB) following the NEB online protocol. D_amino_ was then purified with a mini-Quick Spin DNA Columns (Roche) to remove unincorporated ATP-γ-32P, followed by annealing of the oligos.

FEN1 with or without PARylation (5 nM) was mixed with the unmodified and dT-C6 modified DNA substrates (10 nM) in 20 µl reaction buffer containing 20 mM Tris-HCl (pH 7.4), 40 mM potassium chloride, 5 mM MgCl2, 0.1 mg/ml BSA and 2 mM DTT. After incubated at room temperature for an indicated period of time, cleavage products (20 µl) were mixed with 20 µl 2 × Formamide gel-loading buffer (10 mM EDTA, 0.025% bromophenol blue, 0.025% Xylene cyanol FF and 0.2% SDS dissolved in formamide), heat denatured at 95°C for 3 min, and separated on a 18% acrylamide gel containing 7 M Urea. Gel was then dried and imaged with a GE Typhoon Phosphorimager.

### Proximity ligation assay (PLA)

Duolink PLA fluorescence assay (Sigma Aldrich, Cat# DUO92101) was performed following manufacturer’s instructions. In brief, U2OS cells were seeded on coverslips and treated with CPT for 30 min. After treatment, cells were washed with 1 × PBS and fixed for 15 mins at 4°C in 4% paraformaldehyde in PBS and permeabilized with 0.25% Triton X-100 in PBS for 15 mins at 4°C. The coverslips were blocked with Duolink blocking solution and incubated with indicated antibodies in the Duolink antibody diluent overnight, followed by incubation with PLUS and MINUS PLA probes, ligation and amplification. Coverslips were then washed and mounted using mounting medium with DAPI. Images were captured on a wide field microscope, processed using ImageJ and analyzed using Imaris.

### Detection of DNA-protein crosslinks and their ADP-ribosylation

After FA treatment, 1 × 10^6^ human cells in 35 mm dish per sample were washed with 1 × PBS and lysed with 600 μl DNAzol (Invitrogen), followed by precipitation with 300 μl 200 proof ethanol. The nucleic acids were collected, washed with 75% ethanol, resuspended in 200 μl TE buffer then heated at 65°C for 15 minutes, followed by shearing with sonication (40% output for 10-sec pulse and 10 sec rest for 4 times). The samples were centrifuged at 15,000 rpm for 5 min at 4 °C and the supernatant were collected. 1 μl of the sample was removed for spectrophotometric measurement of absorbance at 260 nm to quantitate DNA content (NanoDrop). 10 μg of DNA from each sample was digested with 50 units micrococcal nuclease (100 units/μl, Thermo Fisher Scientific) in presence of 5 mM CaCl_2_, followed by gel electrophoresis on 4-15% precast polyacrylamide gel (Bio-Rad) for detection of total DPCs using Coomassie stain or silver stain and PARylated DPCs using anti-PAR antibody. Due to the extremely low abundance of PARylated DPCs, samples were run in parallel gels to detect total and PARylated DPCs separately instead of stripping and reprobing the same membrane for their detection. In addition, 2 μg of each sample was subjected to slot-blot for immunoblotting with anti-dsDNA antibody as a loading control to verify that amounts of DNA were digested with micrococcal nuclease.

### *In vivo* complex of enzyme (ICE) assay and mass spectrometry profiling

Cells were exposed to 400 µM for 2 hours and then lysed in sarkosyl solution (1% w/v). The lysates were sheared with a 25-gauge 5/8 needle and loaded onto CsCl solution (150% w/v) for ultracentrifugation in NVT 65.2 rotor (Beckman coulter) at 42,000 rpm for 20 hours at 4°C. The resulting nucleic acid pellets were retrieved and suspended in TE buffer. For mass spectrometric analysis, ICE samples were treated with ribonucleases A and T1 (RNase A/T1) mix to eliminate RNA contamination, followed by addition of 1/10 volume of 3 M sodium acetate sodium acetate and 2.5 volume of 200 proof ethanol. After 20 mins full speed centrifugation, the resulting DPC-containing DNA samples were retrieved, resuspended in ddH_2_O, and digested with micrococcal nuclease (100 units per sample) for 1 h at RT to release the crosslinked proteins. Released protein samples were in-solution digested with trypsin following the filter-aided sample preparation protocol. Dried peptides were solubilized in 2% acetonitrile, 0.5% acetic acid, and 97.5% water for mass spectrometry analysis. They were trapped on a trapping column and separated on a 75 μm by 15 cm, 2-μm Acclaim PepMap reverse phase column (Thermo Fisher Scientific) using an UltiMate 3000 RSLCnano HPLC (Thermo Fisher Scientific). Peptides were separated at a flow rate of 300 nl/min followed by online analysis by tandem mass spectrometry using a Thermo Orbitrap Fusion mass spectrometer. Peptides were eluted into the mass spectrometer using a linear gradient from 96% mobile phase A (0.1% formic acid in water) to 55% mobile phase B (0.1% formic acid in acetonitrile). Parent full-scan mass spectra were collected in the Orbitrap mass analyzer set to acquire data at 120,000 full width at half maximum resolution; ions were then isolated in the quadrupole mass filter, fragmented within the HCD (higher-energy collisional dissociation) cell (HCD normalized energy of 32%; stepped, ±3%), and the product ions were analyzed in the ion trap. Proteome Discoverer 2.2 (Thermo Fisher Scientific) was used to search the data against human proteins from the UniProt database using SequestHT. The search was limited to tryptic peptides, with maximally two missed cleavages allowed. Cysteine carbamidomethylation was set as a fixed modification, and methionine oxidation was set as a variable modification. Diglycine modification to lysine was set as a variable modification for experiments to identify sites of enzymatic PTMs. The precursor mass tolerance was 10 parts per million, and the fragment mass tolerance was 0.6 Da. The Percolator node was used to score and rank peptide matches using a 1% false discovery rate.

### Detection and quantitation of 8-OXO-dG and AP sites

DNA form HEK293 cells were prepared using RADAR assay following treatment with agents FA, campothecin or doxorubicin. Detection and quantitation of 8-OXO-2’-deoxyguanosine (8-OXO-dG) and apurinic or apyrimidinic (AP or abasic) sites in the samples were performed using HT 8-oxo-dG ELISA Kit (R&D Systems) and AP Sites Quantitation Kit (Cell Biolabs), respectively, following manufacture’s instruction. Analyses were conducted using GraphPad Prism.

### FLAG immunoprecipitation (IP)

Cells were washed with 1 × PBS and lysed in 200 μl IP lysis buffer (5 mM Tris-HCl pH 7.4, 150 mM NaCl, 1 mM EDTA, 1% NP-40, 0.2% Triton X-100, 5% glycerol, 1 mM DTT, 20 mM N-ethylmaleimide (Sigma Aldrich) and protease inhibitor cocktail) on a shaker for 15 min at 4 °C, followed by sonication and centrifugation. The supernatant was collected and treated with 1 μl benzonase (250 units/μl, EMD Millipore) for 1h. An aliquot (20 μl) of the lysate of each treatment group was saved as input. Lysates were resuspended in 800 μl IP lysis buffer containing 2.5 μl anti-FLAG M2 antibody (Sigma Aldrich) and rotated overnight at 4°C. 50 μl Protein A/G PLUS-agarose (Santa Cruz Biotechnology) slurry was added and incubated with the lysates for another 4 hrs. After centrifugation, immunoprecipitates were washed with RIPA buffer 2 times then resuspended in 2 × Laemmli buffer for SDS-PAGE and immunoblotting with various antibodies as indicated.

### Molecular combing assay

Fork stability was analyzed using a molecular combing assay. Briefly, U2OS cells were labeled with CldU (100 μM) for 1h, and the cells were sequentially labeled with IdU (100 μM) together with treatment with formaldehyde (400 μM) and FEN1i (10 μM), and in combination for 4h. Next, cells were trypsinized and mixed 1.5% low melting agarose, and then solidified as DNA plugs. The plugs were incubated in lysis buffer (1 mg/ml Proteinase K, 100 mM EDTA, 1% sarkosyl, 10 mM Tris-Cl pH 8.0) overnight at 50 ‘c. The next day, the plugs were gently washed with TE buffer and transferred into 1.6ml of 0.1 M MES (pH 6.5) to melt agarose for 20 min at 70 °C. The agarose was destructed by incubation with β-agarase (NEB) overnight at 42’c and transferred to slides to stretch DNA fibers. Slides were dried for 2 h at 60’c and incubated with 0.4 M NaOH for 30 min. Slides were dehydrated and sequentially incubated with anti-mouse BrdU antibody (347580; BD Biosciences) and anti-rat BrdU antibody (ab6326; Abcam) overnight at 4’c. Following incubation with goat anti-rat secondary antibody (cy5; ab6565; Abcam) and goat anti-mouse secondary antibody (cy3; ab6946; Abcam) for 2 h, slides were incubated with anti-mouse ssDNA primary antibody (MAB3034, Millipore) and secondary antibody BV480 (115-685-166; Jackson ImmunoResearch). Slides were mounted and scanned by FiberVison Automated Scanner (Genomic Vision). The relative ratio between CldU fibers and IdU fibers was calculated from captured images (NImage50/each) and plotted using GraphPad Prism version 9.2.0.

### Flow cytometry with EdU and DAPI

Replication activity was examined by using Click-iT™ Plus EdU Flow Cytometry Assay Kits according to the manufacturer’s instructions (C10634; ThermoFisher Scientific). Briefly, cells were labeled with EdU (10 μM) for 30 min prior to cell harvest. Cells then were fixed with 100 μL of Click-iT™ fixative (Component D) and resuspended the cells in 100 μL of 1X Click-iT™ saponin-based permeabilization reagent. After the washing steps, fluorescent signals were confirmed by flow cytometry using a FACS Canto (Becton Dickinson). Data were analyzed using FlowJo™ Software.

### Neutral comet assay

The comet assays were performed according to the Trevigen CometAssay™ kit protocol with slight modifications. Cells were pre-treated with PARGi for 1 h, followed by co-treatment with 20 μM CPT for 2 h. Treated cells were trypsinized at 37 °C for 5 min. Equal amount of drug-free medium was then added to quench the trypsin activity. The cells were spun down and resuspended in fresh PBS. The final cell density was approximately 100,000 cells/ml. 50 μl of the cell suspension was then mixed with 500 μl of 0.5% low melting point agarose (Invitrogen) (in PBS) at 37 °C. 50 μl of the cell/agarose mixture was transferred onto glass slides. The slides were then immersed in prechilled lysis buffer (2.5 M NaCl, 100 mM EDTA, 10 mM Tris, pH 10.0, 1% Triton X-100, and 10% Me_2_SO) for 1 h. For alkaline comet assay, the slides were immersed in alkaline unwinding solution (200 mM NaOH, 1 mM EDTA) for 30 min at room temperature, followed by electrophoresis in 4°C alkaline electrophoresis solution (300 mM NaOH, 1 mM EDTA) at 1 volt/cm for 30 min. For neutral comet assay, the slides were immersed in 1 X TBE buffer for electrophoresis at 1 volt/cm for 30 min at room temperature. For both alkaline and neutral comet assays, the slides were immersed in 70 % EtOH for 5 min after electrophoresis then incubated with SYBR® Gold for 30 min. The images were visualized under BioTek Cytation 5 cell imaging reader. Statistical analysis was performed by OpenComet, an ImageJ plugin.

### Instant structural illumination microscope (iSIM)

U2OS cells were transfected with FEN1-GFP and TOP1-HaloTag expression plasmids and exposed to FA (400 μM) for indicated time points following 48 h transfection. Live cells were imaged under a customized instant structural illumination microscope. For γH2AX immunofluorescence, 2 mM thymidine was added to U2OS cells in chamber slides at 37 °C for 18 h. Thymidine was removed and fresh DMEM medium was added to the slides for incubation for 9 h. 2 mM thymidine was added to cells for another 18 h incubation. Cells are released from G1/S boundary by washing with PBS and incubating in fresh medium. PARGi (10 μM) and IdU (100 μM) were added to cells 1 h before FA (400 μM) treatment. Cells were collected at indicated time points upon exposure to CPT and washed with PBS, followed by fixation with 4% PFA for 15 min at room temperature. Cells were then incubated with 1.5 M HCl for 30 min at room temperature for DNA denaturation, followed by permeabilization with 0.25% Triton X-100 in PBS (PBST). Cells were blocked with 1% BSA in 0.1% PBST for 30 min, followed by incubation with rabbit anti-γH2AX antibody and mouse anti-BrdU antibody overnight at 4 °C. The next day, Alexa Fluor 568-conjugated anti-rabbit 2^nd^ antibody and Alexa Fluor 488-conjugated anti-mouse antibody were added to chamber slide for 1 h at room temperature. The slides were incubated with DAPI and mounted using ProLong™ antifade mountant. Statistical analysis was performed by ImageJ software.

### Micronucleus analysis

MCF7 WT and shFEN1 cells seeded in chamber slide were exposed to either DMSO or 500 μM FA for 2 h at 37°C. Cells were collected at indicated time points and washed with PBS, followed by fixation with 4% PFA for 15 min at room temperature. Chamber slides were mounted with DAPI, and interphase cells and micronuclei were measured with Nikon SoRa super-resolution spinning disk microscope.

### Processing of PARylated protein samples for mass spectrometry

Agarose beads containing immunoprecipitated proteins were centrifuged at 1500 x g for 1 min (room temperature) and supernatants were removed. The beads were suspended with 25 μl of S-Trap lysis buffer (5% SDS, 50 mM TEAB pH 8.5) and reduced with 20 mM Tris (2-carboxyethyl) phosphine (TCEP) at 56^0^C for 1 h. The samples were alkylated with 20 mM iodoacetamide at room temperature for 30 minutes in the dark. The samples were acidified with 2.5 μl of 27.5% phosphoric acid to completely denature the proteins. The samples were mixed with 170 μl of S-Trap binding buffer and loaded onto S-Trap columns (PROTIFI). The columns were centrifuged at 4000 x g for 30 secs to allow the colloidal proteins to trap in the columns. The columns were washed four times with 200 μl of S-Trap binding buffer. The columns were again centrifuged at 4000 x g for 1 min to remove any residual buffers. Following washing, the trapped proteins in the columns were digested with 2 μg of trypsin suspended in 50 mM TEAB at 37^0^C for overnight. Next day the tryptic peptides were sequentially eluted with 40 μl of 50 mM TEAB (pH 8.5), 0.2% formic acid, and 50% acetonitrile. The eluted samples were lyophilized and subjected to PAR modification removal as described below.

### Removal of PAR modification from tryptic peptides

The lyophilized peptides were suspended in PDE buffer (100 mM HEPES pH 8.0, 15 mM MgCl2). Subsequently, 25 μg of peptides from each sample was treated with 4 μg of NudT16 enzyme at 37^0^C for 2 h to remove PAR groups attached to the peptides. The treated peptides were finally desalted using C18 columns followed by LC-MS/MS mass spectrometry.

### Statistical Analyses

All experiments were repeated at least 3 independent times otherwise stated in the figure legend. Error bars on bar graphs represent standard deviation (SD) except where stated otherwise. The p-value was calculated using paired student’s t-test for independent samples. Statistical analyses were performed using GraphPad Prism software. The differences with *p < 0.05, **p < 0.01 or ***p < 0.001 were considered statistically significant.

