## Supplementary Information for "Flap endonuclease 1 repairs DNA-protein crosslinks via ADP-ribosylation"

**Supplementary Figure 1.**

The common FA-induced protein adducts across the three cell lines was mapped to the human proteome, which was annotated with Gene Ontology (GO) biological process. Enrichment analysis was performed by fold change measurement.

**Supplementary Figure 2.**

**A.** Western blotting in HEK293 cells confirming siRNA knockdown of genes tested by the modified RADAR assay in Fig. 2C by their respective antibodies. **B.** Western blotting in MCF7 cells confirming knockdown of FEN1. **C.** Viability curve derived from ATPlite luminescence assay MCF WT and shFEN1 cells treated with FA at indicated concentrations for 72 h (mean ± SD, n = 3). **D.** The modified RADAR assay was performed in MCF7 WT and shFEN1-transfected cells treated with 400 μM FA for indicated periods of time. Total DPCs were detected with Coomassie stain. **E.** Densitometric analysis comparing total DPC signals generated from the modified RADAR assays including blot shown in Fig. 2D. Density of total DPCs/density of DNA of each group was normalized to cells treated with FA alone. n = 3 independent experiments. Data are presented as mean ± SD. *, $p < 0.05$.

**Supplementary Figure 3.**

**A.** Western blotting in HEK293 cells confirming siRNA knockdown of OGG. **B.** Densitometric analysis comparing total DPC signals generated from the modified RADAR assays including blot shown in Fig. 3D. Density of total DPCs/density of DNA of each group was normalized to cells transfected with control siRNA (siCtrl). n = 3 independent experiments. Data are presented as mean ± SD. NS, not significant. **C.** Quantitation of PLA foci indicating TOP1 and 8-OXO-dG interaction with mean ± SD using Thunderstorm, a plugin of ImageJ. Data were obtained from experiments shown in Fig. 3E. n = 200 biologically independent cells.

**Supplementary Figure 4.**

**A.** IdU/CIdU Ratio with mean ± SD measured from experiments shown in Fig. 4B. ***, $p < 0.001$. **B.** EdU incorporation was analyzed by flow cytometry. Cells were treated with FEN1i at indicated concentrations for 4 h and pulsed with EdU (10 μM) for 30 min prior to harvesting. **C.** Quantitation of tail moments from experiments shown in Fig. 4D using OpenComet, a plugin of ImageJ. n = 200 biologically independent cells. **, $p < 0.01$. **D.** Quantitation of γH2AX intensity from experiments shown in Fig. 4E using ImageJ. n = 200 biologically independent cells. ***, $p < 0.001$. **E.** Quantification of micronucleus containing interphase cells (n > 300 biologically independent cells, error bars = SD) from experiments shown in Fig. 4F.

**Supplementary Figure 5.**

**A.** DNA slot-blot measured by the ICE assay in Fig. 5A. **B.** Densitometric analysis comparing TOP2α-DPC signals generated from the RADAR assays including blot shown in Fig. 5C. Density of TOP2α-DPCs/density of DNA of each group was normalized to cells treated with FA alone. n = 3 independent experiments. Data are presented as mean ± SD. **C.** Densitometric analysis comparing TOP2α-DPC signals generated from the RADAR assays including blot shown in Fig. 5D. Density of TOP2α-DPCs /density of DNA of each group was normalized to cells transfected with control siRNA (siCtrl). n = 3 independent experiments. Data are presented as mean ± SD.

**Supplementary Figure 6.**

**A.** HEK293 cells pre-treated with 10 µM PARGi for 1 h then co-treated with 400 µM FA for indicated periods of time, followed by the modified RADAR assay to detect the kinetics of DPC PARylation using an anti-PAR antibody. **B.** HEK293 cells pre-treated with 10 µM PARGi for 1h and then co-treated with 400 µM FA for 2 h, followed by the modified RADAR assay. Instead of micrococcal nuclease, the RADAR samples were digested with or without recombinant FEN1 in FEN1 cleavage buffer before SDS-PAGE electrophoresis. Total DPCs were detected by Coomassie stain and PARylated DPCs were probed with anti-PAR antibody. *, BSA. **C.** Densitometric analysis comparing total DPC signals generated from the modified RADAR assays including blot shown in Fig. 6C. Density of total DPCs/density of DNA of each group was normalized to cells treated with FA alone. n = 3 independent experiments. Data are presented as mean ± SD. **D.** Densitometric analysis comparing TOP2α-DPC signals generated from the RADAR assays including blot shown in Fig. 6D. Density of TOP2α-DPCs/density of DNA of each group was normalized to cells treated with ETOP alone. n = 3 independent experiments. Data are presented as mean ± SD. **E.** Quantitation of PLA foci indicating TOP1-FEN1 interaction with mean ± SD using Thunderstorm. Data were obtained from experiments shown in Fig. 6E. n = 200 biologically independent cells. **F.** Quantitation of PLA foci indicating TOP2α-FEN1 interaction with mean ± SD using Thunderstorm. Data were obtained from experiments shown in Fig. 6F. n = 200 biologically independent cells.

**Supplementary Figure 7**

**A.** PARylated proteins identified in Fig. 7A were subjected to network analysis using the STRING database, at the interaction confidence setting of 0.9. Proteins not connected to the network were omitted. FEN1 is highlighted in yellow. **B.** Following transfection of empty vector (EV) or FEN1-FLAG expression plasmid, HEK293 cells were treated with the indicated drugs (10 µM PARGi pre-treatment for 1 h followed by 1 h co-treatment with 400 µM FA) and subjected to IP using anti-FLAG antibody. The immunoprecipitates and input samples were Western blotted with the indicated antibodies. **C.** Quantitation of PLA foci indicating TOP1-FEN1 interaction with mean ± SD using Thunderstorm. Data were obtained from experiments shown in Fig. 7E. n = 200 biologically independent cells. ***, $p < 0.001$. **D.** Quantitation of PLA foci indicating TOP2α-FEN1 interaction with mean ± SD using Thunderstorm. Data were obtained from experiments shown in Fig. 7F. n = 200 biologically independent cells. ***, $p < 0.001$. **E.** Densitometric analysis comparing total DPC signals generated from the modified RADAR assays including blot shown in Fig. 7G. Density of total DPCs/density of DNA of each group was normalized to cells treated with FA but without FEN1-FLAG expression plasmid transfection (empty vector, EV). n = 3 independent experiments. Data are presented as mean ± SD. *, $p < 0.05$. **E.** Densitometric analysis comparing TOP2α-DPCs generated from the RADAR assays including blot shown in Fig. 7H. Density of TOP2α-DPC/density of DNA of each group was normalized to cells treated with ETOP but without FEN1-FLAG expression plasmid transfection (empty vector, EV). n = 3 independent experiments. Data are presented as mean ± SD.

**Supplementary Figure 8.**

Working models for the role of PARylation in non-enzymatic and enzymatic DPC repair by FEN1. **A.** DPCs formed within the 5'-flap of Okazaki fragments during strand displacement are readily sensed by PARP1 leading to PARylation, which in turn activates FEN1 for repair following prompt dePARylation of the DPCs by PARG. Concurrent damaged bases adjacent to non-end DPCs (in the front of the fork and on the leading strand behind the fork) are converted into DPC-harboring 5'-flaps by the BER proteins with single-strand breaks that can be detected by PARP1 for PARylation to signal FEN1. FEN1 enriched near Okazaki fragments are in PARylated by PARP1 for their translocation to the front of fork and to the leading strand behind the fork to cleave DPC-harboring 5'-flaps following dePARylation by PARG. **B.** TOP2α are homodimeric enzymes that can act in front of the replication fork to relax accumulated positive supercoils. The homodimers cut both strands of DNA and then pass another DNA molecule through the break. The 5′ end of each break is covalently linked to the tyrosine in the active center of each of the two subunits of the protein (TOP2αcc). In this configuration, the two sides of the nicked DNA are held together by the strong protein-protein interactions between the two subunits of Top2, allowing the nicks to be faithfully resealed in situ. These transient enzyme-DNA intermediate can be trapped by their inhibitors such as etoposide and doxorubicin. Binding of the inhibitor to the interface between one of the subunits and DNA leads to the formation of a single-stranded 5' TOP2-DPC, which can be converted to a 5' flap structure via post-translational modifications, conformational changes, or polymerase activities. PARP1 senses the TOP2-linked SSB and PARylates the DPC to signal FEN1. In the meantime, FEN1 at Okazaki fragments at the rear of the fork is also PARylated by PARP1, which drives FEN1 to the DPC site for cleavage upon dePARylation by PARG.

### Supplementary figure 1

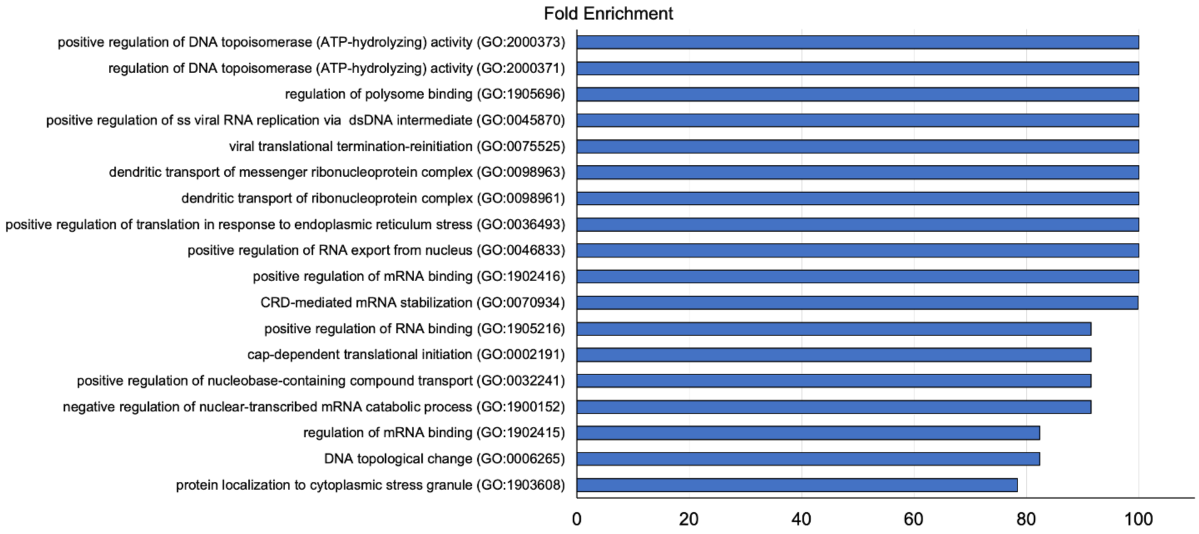

**Supplementary figure 2**

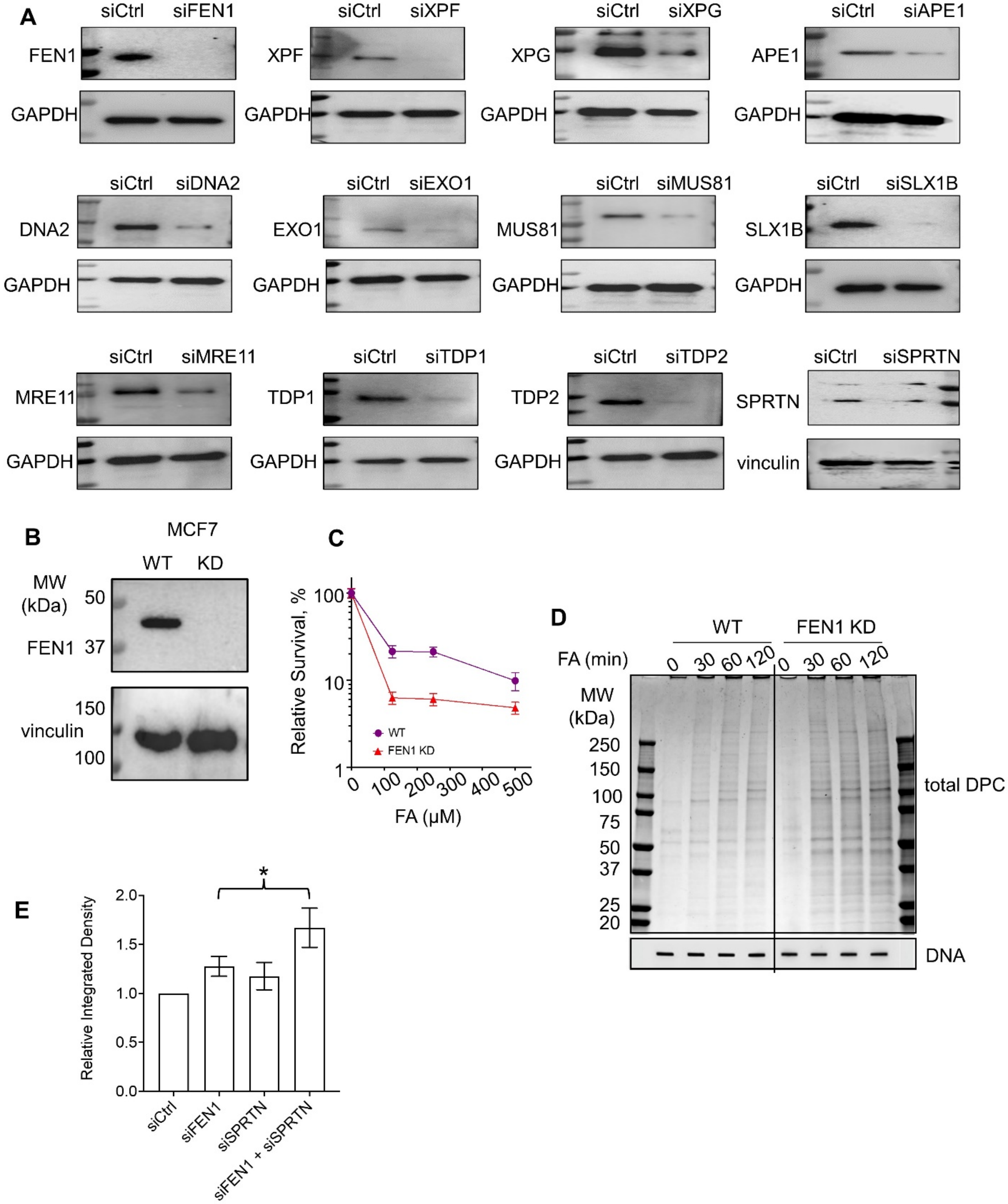

**Supplementary figure 3**

A

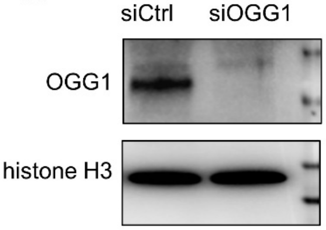


B

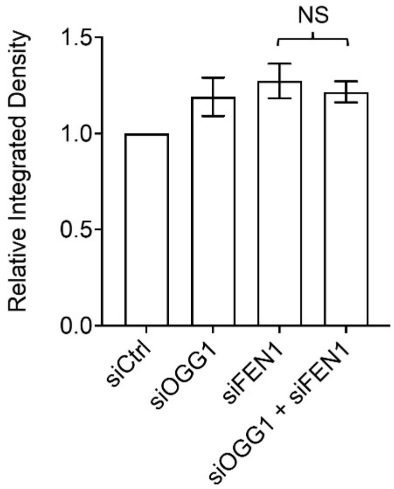


C

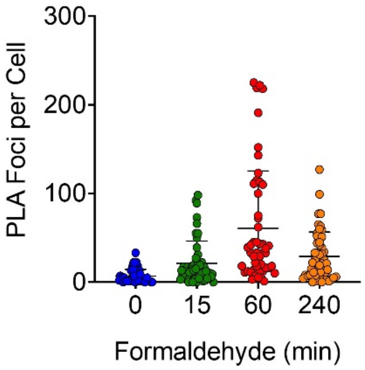

**Supplementary figure 4**

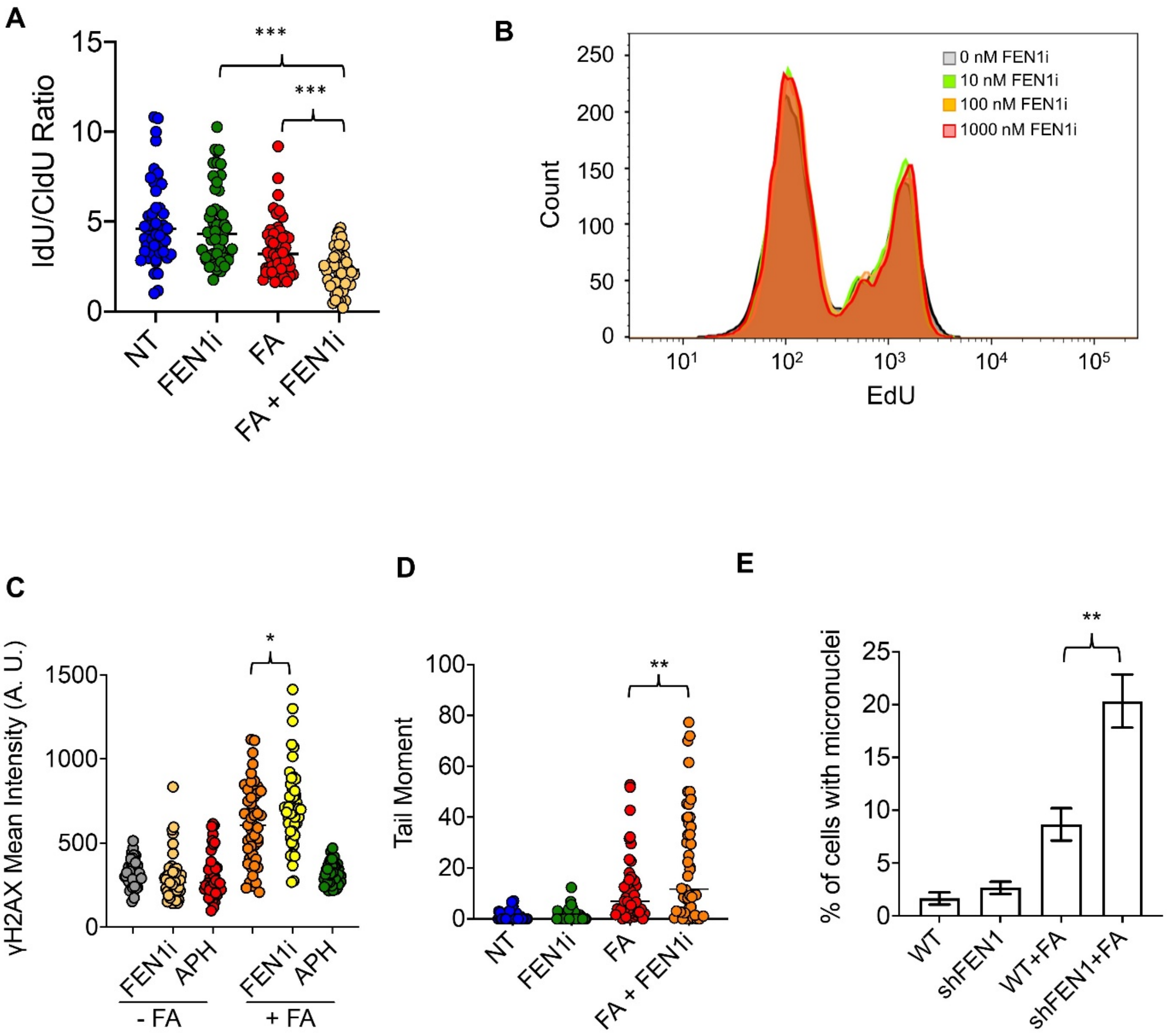

**Supplementary figure 5**

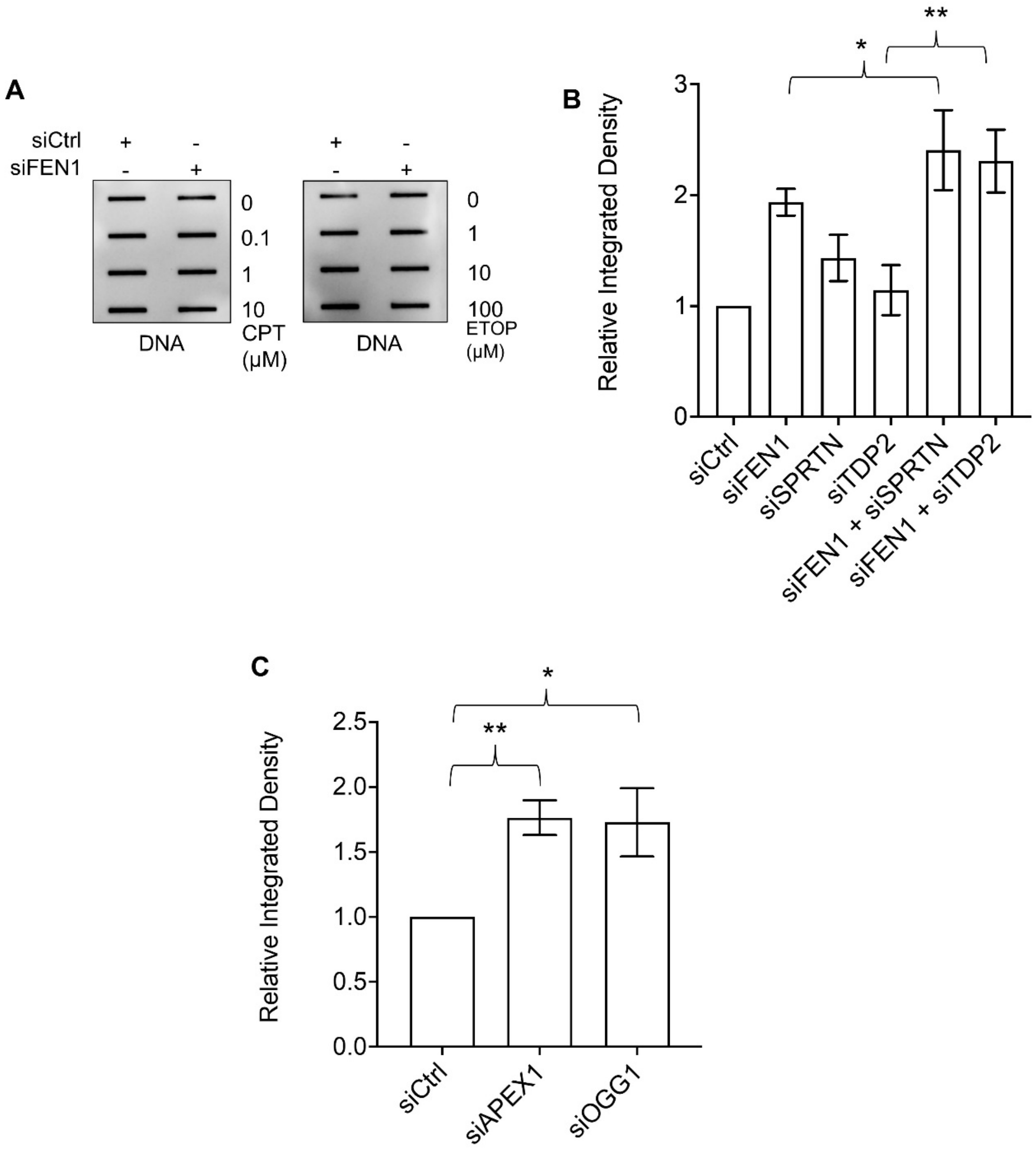

### Supplementary figure 6

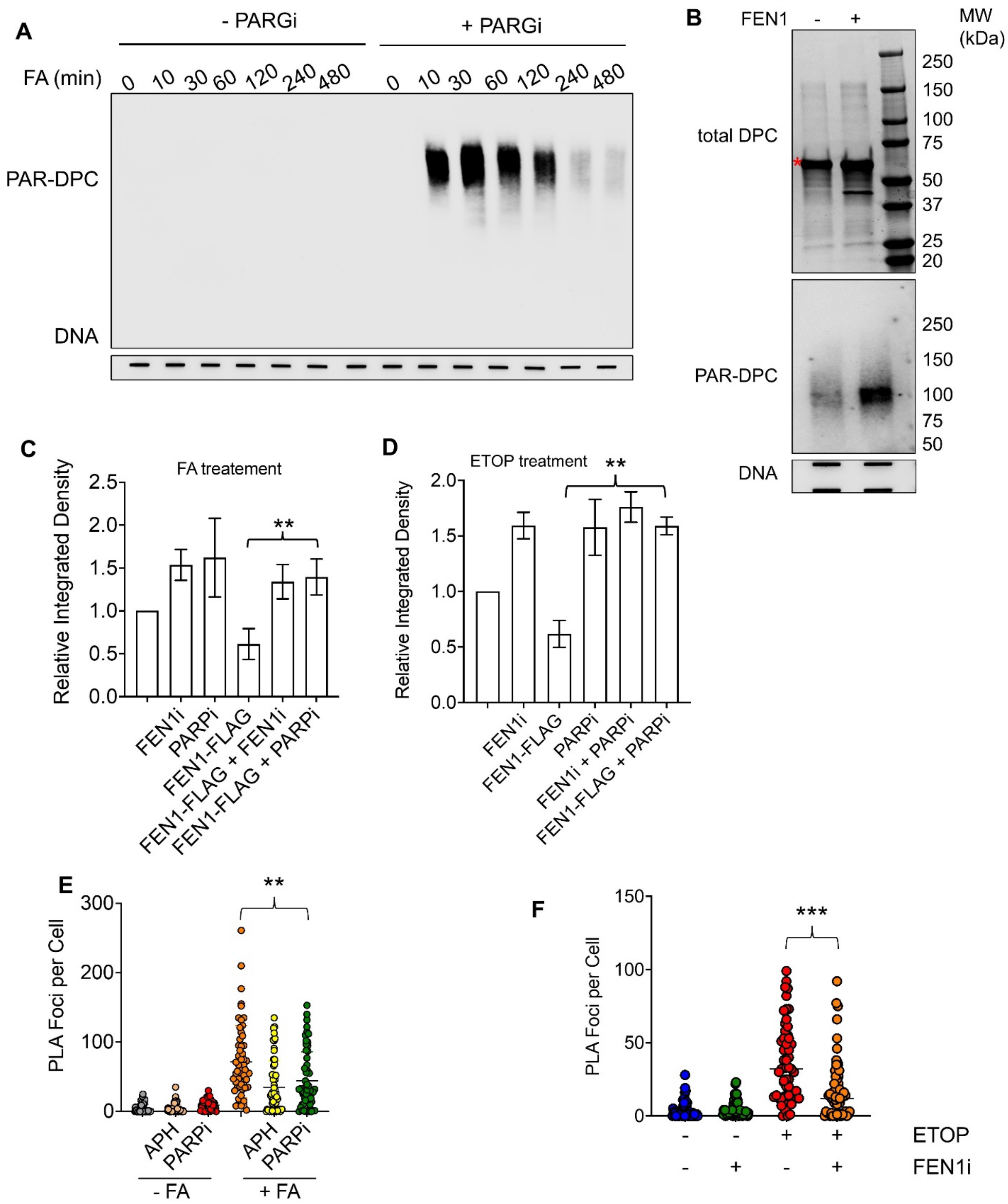

### Supplementary figure 7

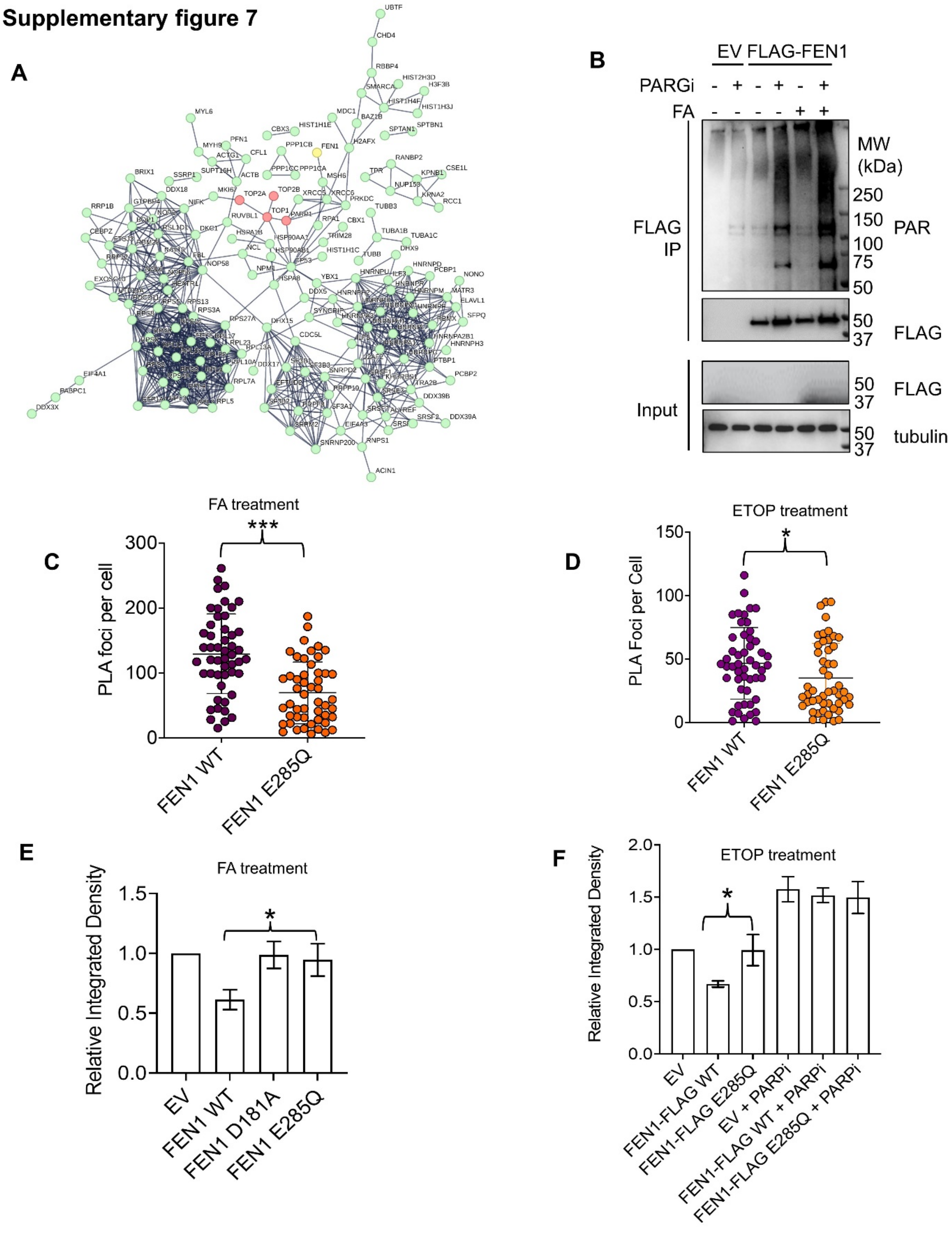

**Supplementary figure 8**

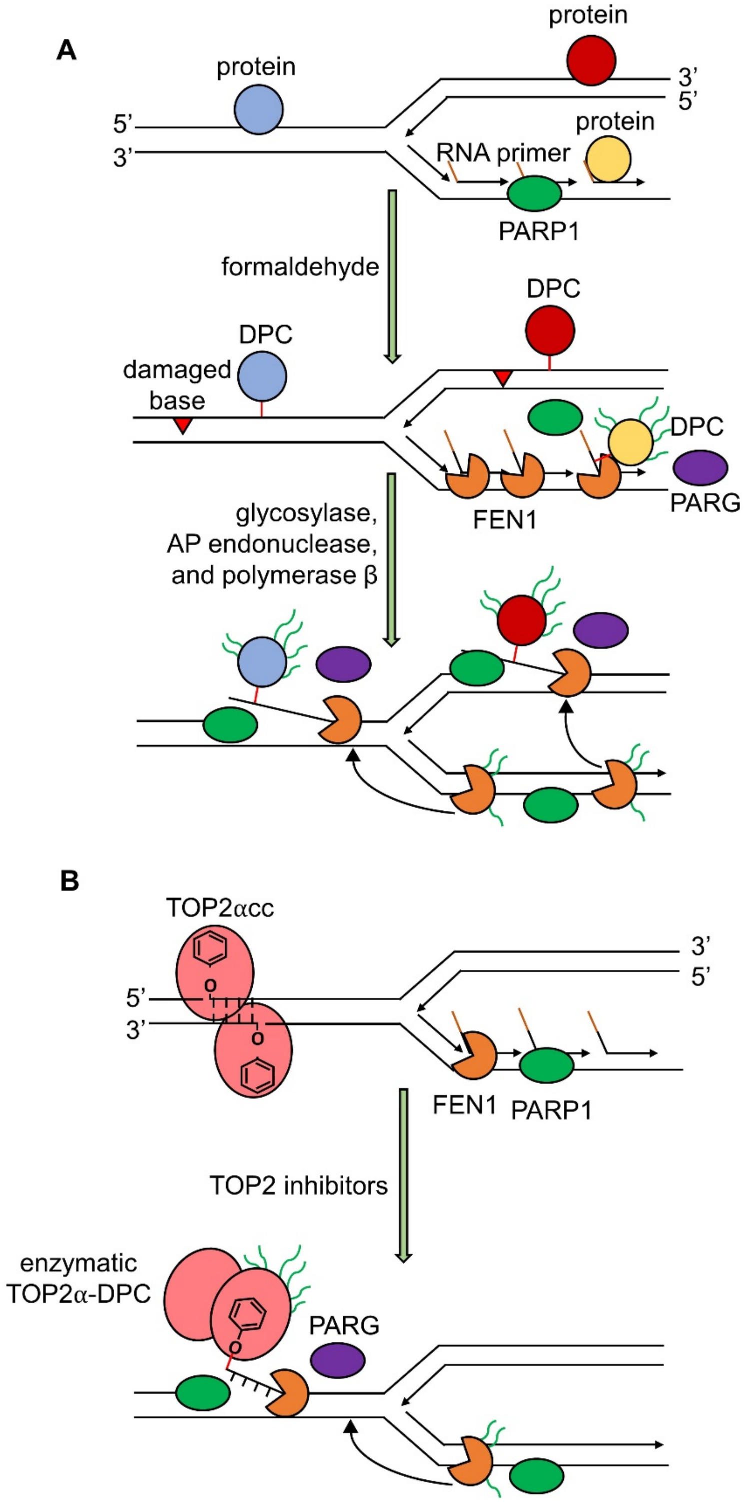
